# Atomic resolution structure of spinach rubisco reveals protons and dynamics

**DOI:** 10.64898/2025.12.22.696010

**Authors:** Nicholas Croy, Guillaume Gaullier, Patricia Saura, Simonas Masiulis, Asmit Bhowmick, Jan Kern, Oliver Raschdorf, Inger Andersson, Ville R. I. Kaila, Cecilia Blikstad, Johannes Messinger

**Author notes:** shared first authors (Nicholas Croy, Guillaume Gaullier). corresponding authors;, **Correspondence and requests for materials** should be addressed to N.C., C.B., or J.M.

## Abstract

Photosynthetic organisms sustain life on Earth by storing solar energy in biomass. Central to this process is rubisco, the enzyme that catalyses the fixation of CO_2_ to ribulose-1,5-bisphosphate, providing the primary gateway for inorganic carbon into the biosphere. Rubisco’s catalytic efficiency is a major determinant of crop productivity and global carbon flux, making it a longstanding target for protein engineering.^1–4^ Yet, attempts to enhance its performance through rational design have met limited success due to an incomplete understanding of rubisco’s catalytic mechanism. Here, we report an atomic resolution (1.25 Å) cryo-EM structure of spinach rubisco in complex with the transition-state analogue 2-carboxyarabinitol-1,5-bisphosphate. Supported by large-scale quantum/classical (QM/MM) calculations, our structural analysis reveals protonation equilibria within the active site and unexpected structural flexibility across large protein regions despite the exceptionally tight ligand binding. Our findings provide new insight into the complex interplay of protonation equilibria and conformational sampling, suggesting a novel basis for rubisco’s rational redesign utilizing strategies that rely on a combination of dynamic and electrostatic control.

## Main

Structural and functional studies have revealed many aspects of how rubisco (ribulose-1,5-bisphosphate carboxylase/oxygenase, EC 4.1.1.39) converts CO_2_ and 5-carbon ribulose-1,5-bisphosphate (RuBP) into two molecules of 3-carbon 3-phosphoglycerate (3PGA) (see reviews ^4–7^). This complex chemical transformation involves a sequence of well-orchestrated enolization, carboxylation, hydration, bond scission, and stereospecific protonation reactions (Fig. 1). Although each of these reaction steps involves at least one proton transfer, the specific active site residues and water molecules responsible for these proton transfers remain contested, giving rise to multiple, competing mechanistic proposals.^8–12^ Moreover, the role of conformational dynamics in promoting carbon fixation is not sufficiently understood. Resolving these mechanistic ambiguities is crucial for further protein engineering efforts aimed at improving rubisco’s catalytic efficiency, which is limited by a low catalytic rate and poor specificity, i.e. the tendency to catalyse a competing oxygenation side-reaction.^13,14^

**Figure 1.**
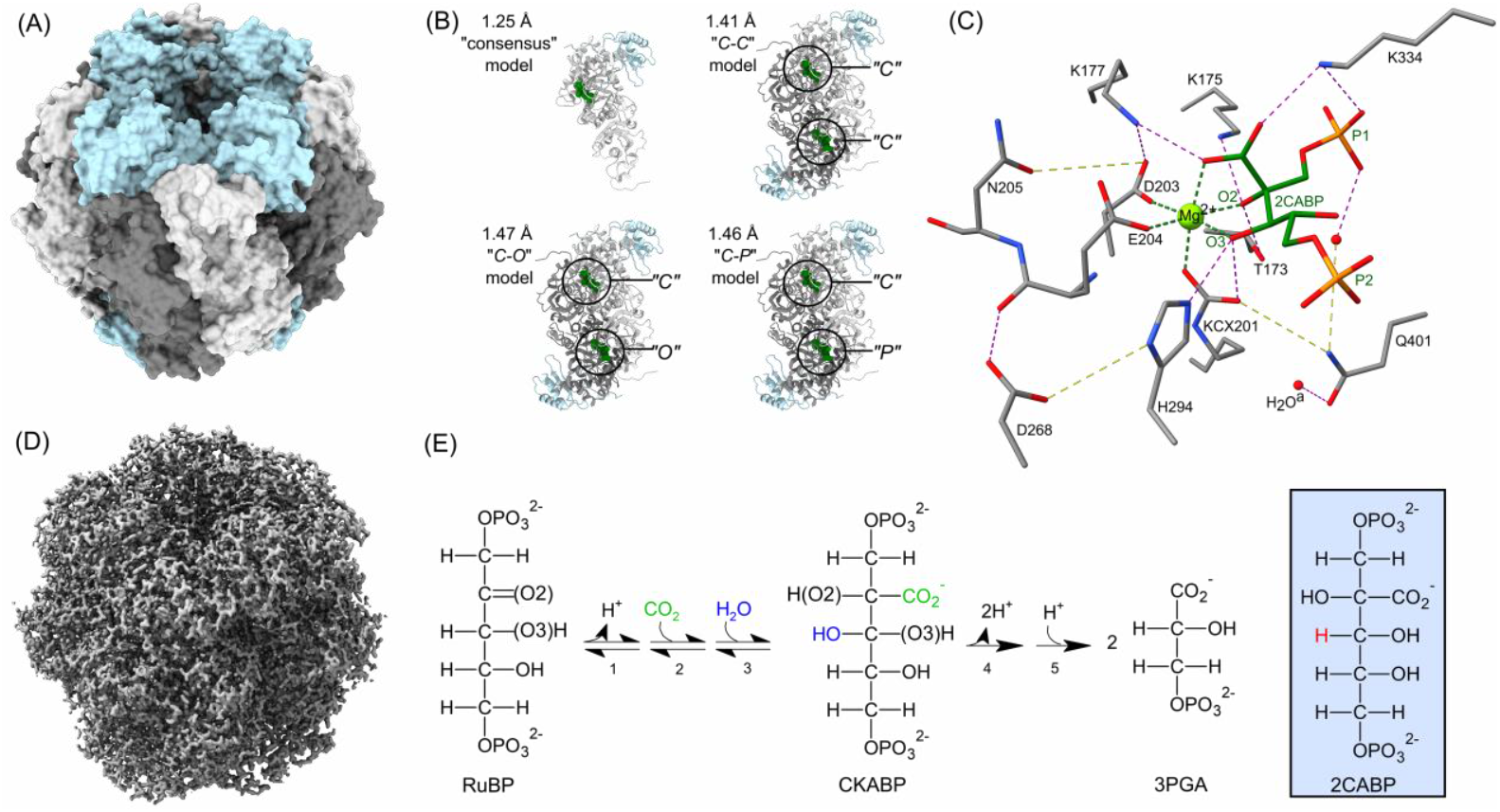
Rubisco’s structure and reaction chemistry. **(A)** The complete L8S8 rubisco assembly. **(B)** The four models obstained from this experiment: one L1S1 consensus model and three L2S2 models. Two active sites are formed at the boundary between two large subunits that form a dimer. The active site-bound transition-state analogue, 2CABP, is shown in dark green. The L2S2 active sites are designated “C”, “P”, or “O” depending on whether the N-domain for that active site is closed, partially open, or open, respectively (see Fig. 3A). **(C)** 2CABP and a subset of key residues, water molecules, and the activating Mg^2+^ ion (light green) in the active site. Green and magenta dashed lines indicate Mg^2+^ coordination and hydrogen bonds, respectively. Yellow dashed lines were added for clarity (distances >4 Å). **(D)** The unsharpened, D4-symmetric consensus reconstruction at level 0.0611 (7σ). **(E)** The carboxylation reaction of RuBP involves (1) enolization, (2) carboxylation, (3) hydration, (4) C2-C3 bond cleavage, and (5) stereospecific protonation. In 2CABP, one of the CKABP C3-OH groups (blue) is replaced with a proton (red). 2CABP resembles the carboxylated reaction intermediate, CKABP.

Rubisco from higher plants assembles into a hexadecameric complex composed of eight large subunits (RbcL, ~55 kDa) and eight small subunits (RbcS, ~18 kDa). The RbcL subunits form head-to-tail dimers, and two active sites are formed at the intra-dimer interface between the carboxy-terminal domain of one RbcL and the amino-terminal domain of a second RbcL (Fig. 1).^7^ However, active rubisco is only formed after carbamylation of an active site lysine residue, K201, with one non-substrate CO_2_ molecule (the resulting carbamylated lysine is hereon referred to as KCX201) followed by the monodentate, facial coordination of one Mg^2+^ ion by KCX201, D203, and E204.^15,16^ The three vacant/water-bound Mg^2+^ coordination sites are then used to coordinate RuBP and its reaction intermediates during the reaction cycle. The O2 and O3 sites of RuBP (Fig. 1E) occupy equatorial positions opposite to D203 and E204, while the substrate CO_2_ likely binds to the axial site opposite to KCX201 after its addition to RuBP.^17^ Figure 1C shows the binding of a transition-state analogue, 2-carboxyarabinitol bisphosphate (2CABP) (Fig. 1C), which resembles the hydrated product after CO_2_ addition, 2-carboxy-3-ketoarabinitol bisphosphate (CKABP). Three residues—KCX201, K175, and H294—are thought to be important for the proton transfers to and from the RuBP-O2 and RuBP-O3 which facilitate rubisco’s reaction chemistry, but their precise roles remain speculative.^8–12^ Here, we aim to provide mechanistic insight by revealing the protonation states of all active site groups, as well as by better resolving structural changes connected to active site opening/closing.

The experimental determination of hydrogen atom positions in protein structures remains a major challenge; each of the various X-ray, neutron, and electron scattering methods used for protein structure determination has unique, demanding requirements. Visualizing hydrogen atoms by X-ray crystallography requires a diffraction resolution of 1.0 Å or higher, which is beyond the reach of most protein crystals. A more modest resolution of 2.5 Å is required to reveal most deuterons by neutron crystallography, but this approach relies on crystal volumes in excess of 0.1 mm^3^, which can be technically difficult to achieve.^18^ For single-particle cryo-electron microscopy (cryo-EM), densities for hydrogen atoms have been reported in structures with resolutions as low as 1.7 Å,^19,20^ while an analysis by Yamashita *et al*. (2021) showed that some hydrogen atoms can be observed even at a 2.0 Å resolution.^21^

Motivated by these findings, we collected cryo-EM data of the activated rubisco in complex with 2CABP (rubisco-Mg^2+^-2CABP complex) and obtained a reconstruction at 1.25 Å resolution, which is at par with the best cryo-EM structures yet obtained.^19,22^ Our data reveal hydrogen atoms throughout the protein (Fig. 2 and Supplementary Fig. 1), including an unexpected active site protonation configuration which is supported by our large-scale hybrid quantum/classical (QM/MM) calculations. Further, we show surprising structural flexibility that includes both alternative conformations of individual side chains as well as fully open, partially open, and closed active site states despite the exceptionally tight (~190 fM)^23^ 2CABP binding. This leads to an expansion of the previous binding model, a reinterpretation of the apparent femtomolar binding constant, and new insights into rubisco’s unique carboxylation mechanism.

**Figure 2.**
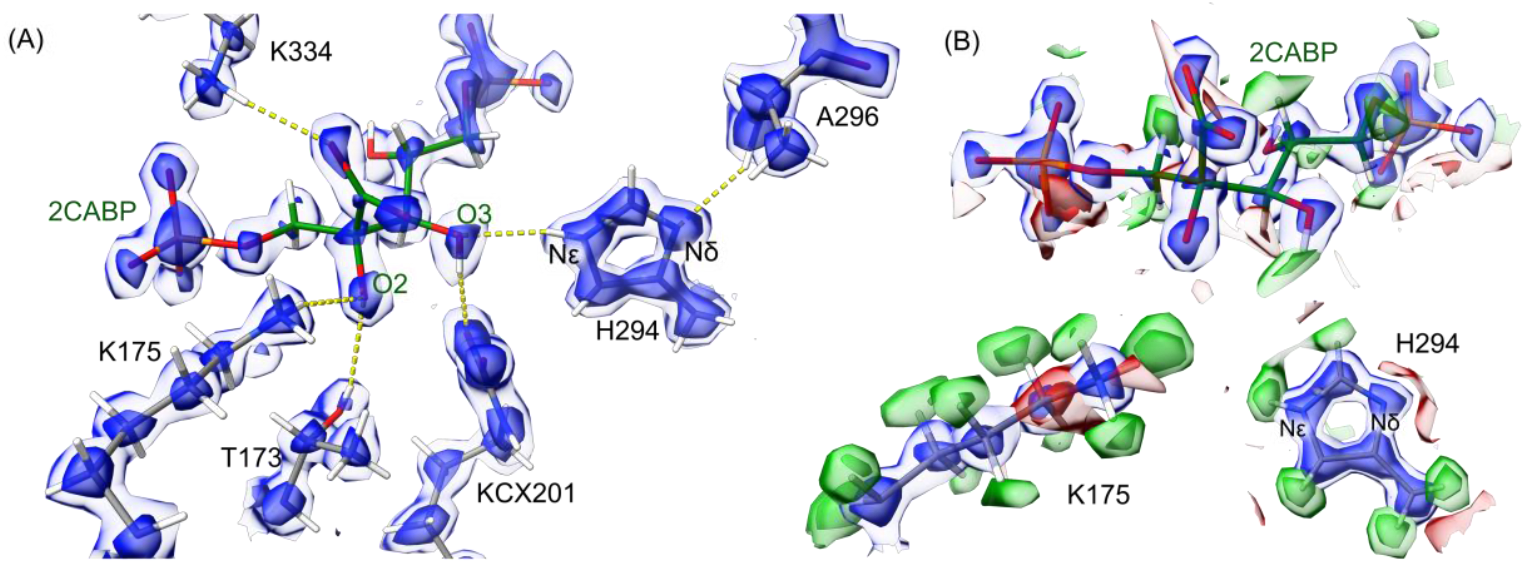
Active site proton configuration. **(A)** The sharpened F_o_ map obtained from Servalcat is shown at two absolute map levels: 1.5 (light blue) and 3.5 (dark blue). Individual C, N, and O atoms are resolved at the higher map levels. Key hydrogen bonds formed between 2CABP and active site residues are indicated by yellow dashed lines. The activating Mg^2+^ is omitted for clarity. **(B)** Both the sharpened F_o_ map (absolute map levels 1.5 and 3.5) and H-omit F_o_-F_c_ difference map are displayed for 2CABP (F_o_-F_c_ absolute map levels 2.0 and 2.75), K175 (F_o_-F_c_ absolute map levels 2.0 and 3.0), and H294 (F_o_-F_c_ absolute map levels 2.0 and 3.5). Different map levels are employed for each of these elements to account for their differing signal strengths resulting from their differing flexibility. The negative red densities are due to this flexibility or negative charges (on the 2CABP), while the positive green densities indicate the position of hydrogen atoms on either the selected side-chain/ligand or neighbouring residues.

## Results

### Atomic-resolution consensus structure

To obtain a high-resolution structure of spinach rubisco, we vitrified the protein on HexAuFoil grids and recorded images on a 300 kV Krios 5 instrument equipped with a cold Field Emission Gun, an Energy Filter, and a Falcon 4i detector operated in electron counting mode, utilizing a pixel size of 0.548 Å and a total dose of 53 electrons per Å^2^. To ensure that our samples were of the highest biochemical quality, all grids were prepared with the protein from one dissolved crystal of the rubisco-Mg^2+^-2CABP complex (see SI for further experimental details). We refined around 1.6 million particles using D4 symmetry to obtain a reconstruction at 1.25 Å resolution (Supplementary Fig. 2). The high-resolution of the present data is apparent throughout the structure, where individual C, N, and O atoms are clearly resolved in the sharpened map at high contour levels (Fig. 2). At lower contour levels, the map shows well-defined additional cryo-EM densities which agree with the positions of hydrogen atoms at almost all expected sites, though the hydrogen atoms are best visualized with an F_o_-F_c_ H-omit difference maps (Fig. 2B-D), where 89% and 85% of the expected hydrogen atoms are detected at 2 and 3 sigma, respectively (see Supplementary Discussion S1 and Supplementary Table 1).

### Conformational Sub-States Revealed by Clustering Analysis

The D4 symmetry imposed during refinement of the consensus reconstruction averages the eight L1S1 units found in one full assembly, improving signal at locations where they are identical, but also dampening the signal from any subpopulations of particles with distinct conformational states that deviate from perfect symmetry. Despite this, the consensus reconstruction reveals weaker additional conformations of the backbone and of amino acid side chains in many areas.

Different conformations of rubisco have been observed previously. X-ray structures capture two distinct states—one in which the active site is “open” and one in which the active site is “closed”— and these X-ray structures show a correlation between the active site state and the ligand inter-phosphate distance (P1-P2 closed < 9.2 Å < P1-P2 open; Figure 1C).^24^ For this reason, rotations of the ligand phosphates that decrease the inter-phosphate distance are thought to trigger active site closure via a cascade of structural changes, including (1) a 2°-rotation of the N-terminal domain toward the active site (“closed N-domain”) and (2) a 12-Å movement of “loop 6” (residues 331-338) into the active site (“closed loop 6”), followed by (3) the ordering of the C-terminal residues (residue 461 to C-terminus) over loop 6 (“closed C-terminus”; Figure 3A).^24,25^

**Figure 3.**
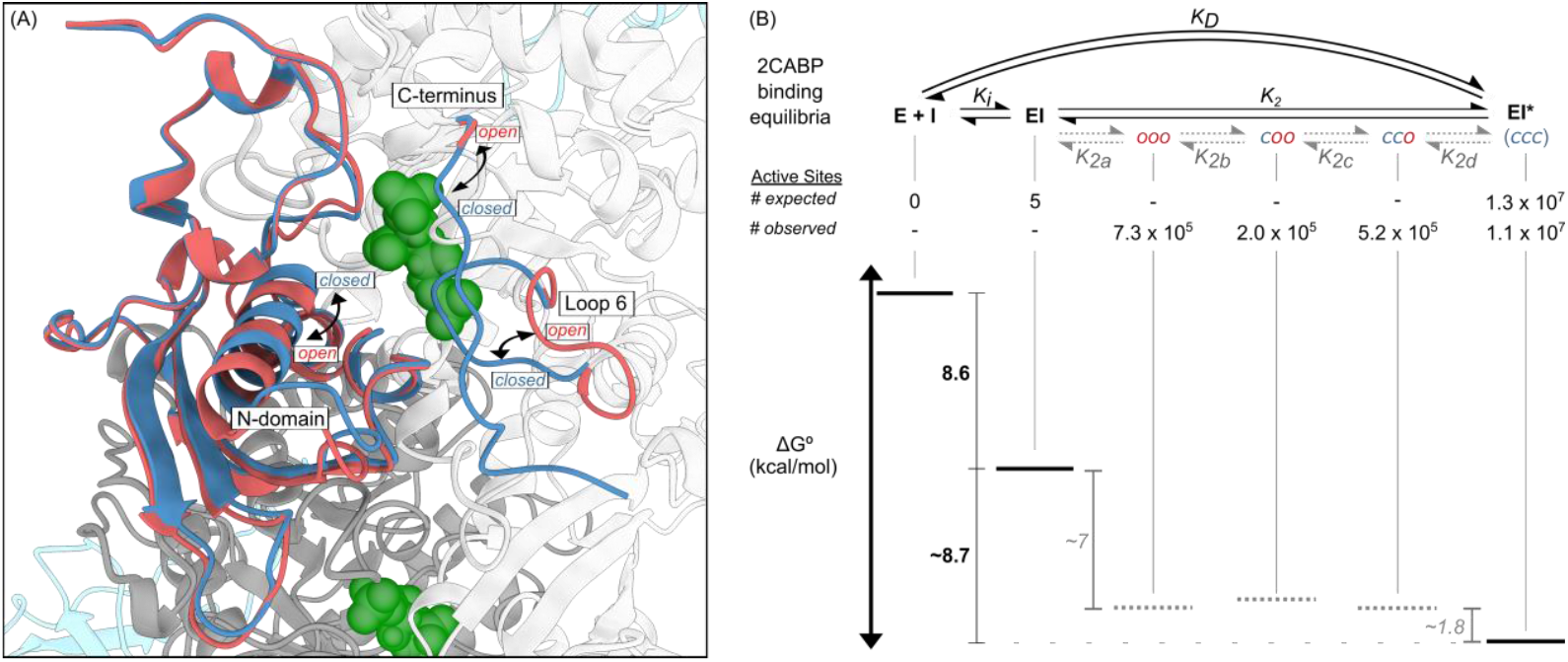
Active site states and their equilibria with 2CABP bound. **(A)** The three primary elements responsible for active site closure, the N-domain, loop 6, and the C-terminus, are shown in their open (red) and closed (blue) positions. As these elements are semi-independent, their combination produces six possible active site states: ooo, coo, cco, ccc, oco, and occ, where lower case c/o represent closed/open states for the N-domain, loop 6, and C-terminus, respectively. **(B) Top:** Previous (black) and proposed (gray) 2CABP binding models. Two states in the proposed binding model, oco and occ, are off-axis and not shown here for simplicity but are shown in Supplementary Discussion S3. **Middle:** The number (#) of expected active sites based on the previous binding model^23,26^ and the # observed active sites in our data. **Bottom:** A free energy diagram based on literature data^23^ and equilibrium constants calculated using the # of active sites found in each state, as estimated by the occupancy of the N-domain, loop 6, and the C-terminus. For the calculations presented we assume that the room temperature distribution of states is preserved during the rapid vitrification. See Supplementary Discussion S2 and Supplementary Table 4 for discussion and summary of assigned states.

For better characterization of the structural heterogeneity in the present rubisco-Mg^2+^-2CABP cryo-EM data, we employed C4 symmetry expansion, which allowed us to create and classify “sub-particles” containing one L2S2 unit with two complete active sites. This approach revealed three types of L2S2 tetramers which, based on the N-domain state in each active site, were classified as *closed - closed* (*C-C*), *closed - partially open* (*C-P*), and *closed - open* (*C-O*), and this single letter code (*C, P, O*) will be used to collectively refer to these groups of sites (Fig. 1B; see Supplementary Table 2 for a summary of the six L2S2 classes). Further analysis evaluating the occupancies of the N-domain, loop 6, and C-terminus (Supplementary Table 3) uncovered a rich structural heterogeneity within each class (SI Text S2, Supplementary Table 4) that required a three-letter nomenclature, where fully closed active sites are denoted “*ccc*” (N-domain, loop 6, and C-terminus in closed positions; Fig. 3A), fully open active sites as “*ooo*” (N-domain, loop 6, and C-terminus in open positions), and intermediate states as e.g. “*coo*” and “*cco*”.

The binding of 2CABP was previously described by a two-step *induced fit* model. In the first step, the inhibitor (I) rapidly binds to the enzyme (E) to form a reversible EI complex (the Michaelis complex). In the second step, induced conformational changes slowly convert EI to a significantly stronger and essentially irreversible EI* state (Fig. 3B, top).^23,26^ Early studies interpreted the slow EI to EI* conversion rate — typical of transition-state analogues — as the inefficient structural rearrangements associated with active site closure, i.e. the slow conversion of the *ooo* state (EI) to the *ccc* state (EI*). However, based on the energetics of this binding model, one would only expect five active sites to populate the *ooo* state in our data, while the remaining 13 million active sites (8 active sites × 1.6 million particles) would populate the *ccc* state. By contrast, our cryo-EM classifications (see SI) reveal 730 000 *ooo*, 200 000 *coo*, 520 000 *cco*, and 11 million *ccc* active sites (Fig. 3B, middle), i.e. about 15% of all active sites are not in the fully closed *ccc* state. This structural diversity likely escaped detection in prior X-ray diffraction studies due to crystal contacts favouring the lowest energy (*ccc*) state, as compared to the vitrification in cryo-EM which preserves the natural conformational diversity of individual particles in solution.

To rationalize these observations, we propose an expanded binding model (Fig. 3B) with free energy levels derived based on the observed populations of the active site states in the presence of 2CABP (see SI Text S3). Our expanded binding model suggests that the exceptional binding affinity of 2CABP originates from processes — the formation of the EI state (8.6 kcal mol^−1^)^26^ and the subsequent formation of the *ooo* state observed here (at about 7 kcal mol^−1^) — occurring *before* active site closure, while active site closure only contributes about 2 kcal mol^−1^.

Previous studies indicate that the carboxyl group of 2CABP (2CABP-CO_2_) is required to form the EI* state^26^, suggesting that interactions with 2CABP-CO_2_ are responsible for the tight binding. In the closed active site, 2CABP-CO_2_ forms stabilizing hydrogen-bonding interactions with K334 (loop 6) and K177 (Fig. 1C). As our data suggests that the loop 6 open/closed states (*coo/cc*o) are nearly isoenergetic, this would indicate that the interaction between 2CABP-CO_2_ and K177 is responsible for the transition from EI to the tightly bound intermediates. However, given that K177 is poised to interact with 2CABP-CO_2_, it is difficult to rationalize the slow onset of EI* formation without invoking additional processes which kinetically gate the transition to EI*, such as, e.g., structural changes extending to the small subunit, which may increase conformational entropy, or prerequisite proton transfers.

### Substrate Water Mobility

The changes in structure associated with active site closure include a reorientation of many sidechains^24^, some of which have been reported based on previous X-ray structures.^9,17^ Here we focus on Q401, which upon active site closure, rotates toward the bound 2CABP. Our data suggest that this rotation displaces an active site water molecule, H_2_O^a^, from its original “proximal” site to a “distal” site, *ca*. 6 Å away (Fig. 4). Due to the short distance between the proximal H_2_O^a^ and the substrate C3 (3.92 Å) in the X-ray structure of the Rubisco-Ca^2+^-RuBP complex, H_2_O^a^ is thought to function as the substrate water molecule in the hydration step (step 3 in the reaction scheme in Fig. 1), though it has been unclear why this proximal water molecule has not been observed in X-ray structures of the closed 2CABP complex.^8–12^

**Figure 4.**
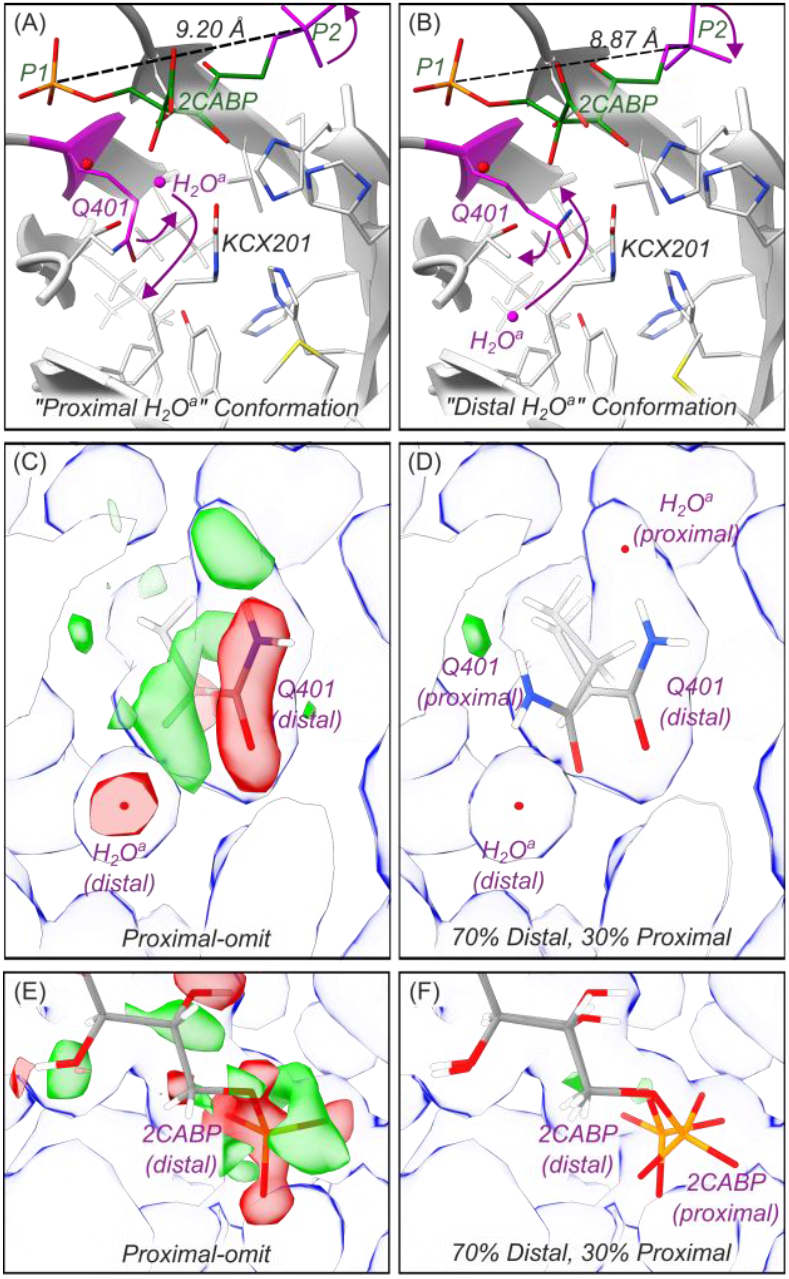
Substrate Water Mobility. **(A)** The “proximal H_2_O^a^” conformation, and **(B)** the “distal H_2_O^a^” conformation are shown from one of the “C” active sites in the C-C model. The proximal position is observed in X-ray structures with an open active site^9^, while the distal conformation is observed in X-ray structures with a closed active site.^17^ Fo-Fc omit maps (absolute levels −1.5, 1.5) generated with a model lacking the proximal positions of Q401 and H_2_O^a^ and 2CABP-P2 are shown in **(C)** and **(E)**, respectively. Positive (green) densities indicate partial occupancy in the proximal positions, while the negative (red) densities indicate that the positions are over-modelled. Modelling the distal and proximal positions with 70% and 30% occupancies, respectively, removes all difference peaks, as shown in **(D)** and **(F)**. Unsharpened maps (blue) are shown at an absolute level of 0.0111 in (C) and (D), and 0.0167 in (E) and (F).

In all our active sites, we see evidence for both Q401/H_2_O^a^ positions, and their occupancies correlate with two 2CABP conformations that differ by a small rotation at 2CABP-P2. We designate the two sets of correlated positions the “proximal H_2_O^a^” and “distal H_2_O^a^” conformations (Fig.4). The occupancy of the “proximal H_2_O^a^” conformation is lowest in the “*C*” active sites, suggesting that a subset of the structural changes required for active site closure is sampled in all active site states, but to different degrees. Further, in the “*C*” active sites, the 2CABP inter-phosphate distance increases on average from 8.87 Å in the “distal H_2_O^a^” conformation to 9.20 Å in the “proximal H_2_O^a^” conformation. Intriguingly, 9.20 Å is the inter-phosphate distance which demarcates the *open* and *closed* active site X-ray models.^24^ Thus, the 2CABP-P2 rotation, which our data show can occur without active site opening, appears to be part of the “subset of structural changes” responsible for modulating the position of H_2_O^a^.

However, it remains unclear if the distal H_2_O^a^ position is functionally relevant, and the described dynamics could rationalize both a stepwise and a concerted mechanism for the carboxylation and hydration steps.^7^ For example, the distal position could serve as a transient stalling site for H_2_O^a^, which moves back into the active site for the hydration reaction after CO_2_ addition in a stepwise mechanism. Alternatively, CO_2_ entry into the active site may induce active site closure, mobilizing H_2_O^a^ for concerted CO_2_ addition and hydration reactions; with 2CABP bound, H_2_O^a^ cannot react in the proximal position, and is thus displaced.

### Active Site Protons

To identify key hydrogen bonding interactions with 2CABP in the active site, we closely analysed the F_o_-F_c_ H-omit difference map of the consensus structure. Several protons involved in 2CABP binding clearly stand out as strong positive difference peaks (Fig. 2B). This reveals that H294 is singly protonated at Nε, thus contradicting previous proposals which assume the protonation of H294 at the Nδ position.^8–11^ Moreover, while H294-Nδ forms a hydrogen bond with the protonated backbone nitrogen of A296, H294-Nε serves as the donor atom in a hydrogen bond formed with 2CABP-O3, which remains protonated and serves as the donor atom in a hydrogen bond formed with KCX201. No protons are visible on 2CABP-O2, which accepts hydrogen bonds from both T173 and a fully protonated K175 (Fig. 2B).

To validate the experimentally assigned protonation states, we performed large-scale hybrid quantum/classical (QM/MM) calculations treating the entire active site – the 2CABP and surrounding first- and second-coordination sphere amino acids (*ca*. 310 atoms) – at the quantum mechanical (density functional theory) level (Supplementary Fig. 5). This enabled us to probe the effect of alternative active site protonation states on both the QM/MM optimised geometries as well as the calculated electrostatic potential (ESP) maps. To this end, we characterised the goodness of fit by (1) calculating the root-mean square deviation (RMSD) of QM/MM-derived models relative to our experimental model, and (2) quantifying the volume of densities in ESP difference maps obtained by subtracting the experimental map from our calculated ESP maps (Supplementary Fig. 6-8, Supplementary Table 6-7). Our QM/MM models strongly support the experimentally assigned protonation states (Fig. 2A), which result in a small RMSD of only 0.149 Å (Supplementary Fig. 6B). In contrast, alternative protonation states for surrounding histidines, including H294, result in significant ESP difference maps, thus further supporting the experimentally assigned states (Supplementary Fig. 6). However, our calculations also indicate that mixed protonation states at 2CABP-O2 and K175 are consistent with the experimental data, suggesting that protonation equilibria are in part responsible for forming the various observed active site states. For example, upon structure relaxation, the [K175^0^, O2-H, O3-H] state, generated by transferring a proton from K175 to 2CABP-O2, is nearly isoenergetic with the experimentally assigned state [K175^+^, O2-, O3-H]. Moreover, our QM/MM calculations suggest that the proton transfer from K175 to 2CABP-O2 results in a rotation of D203, and our calculated QM/MM-ESP map, assuming 50% occupancy for each state, reproduces the experimental ESP asymmetry around D203 (Fig. 5A, B, Supplementary Fig. 8).

**Figure 5.**
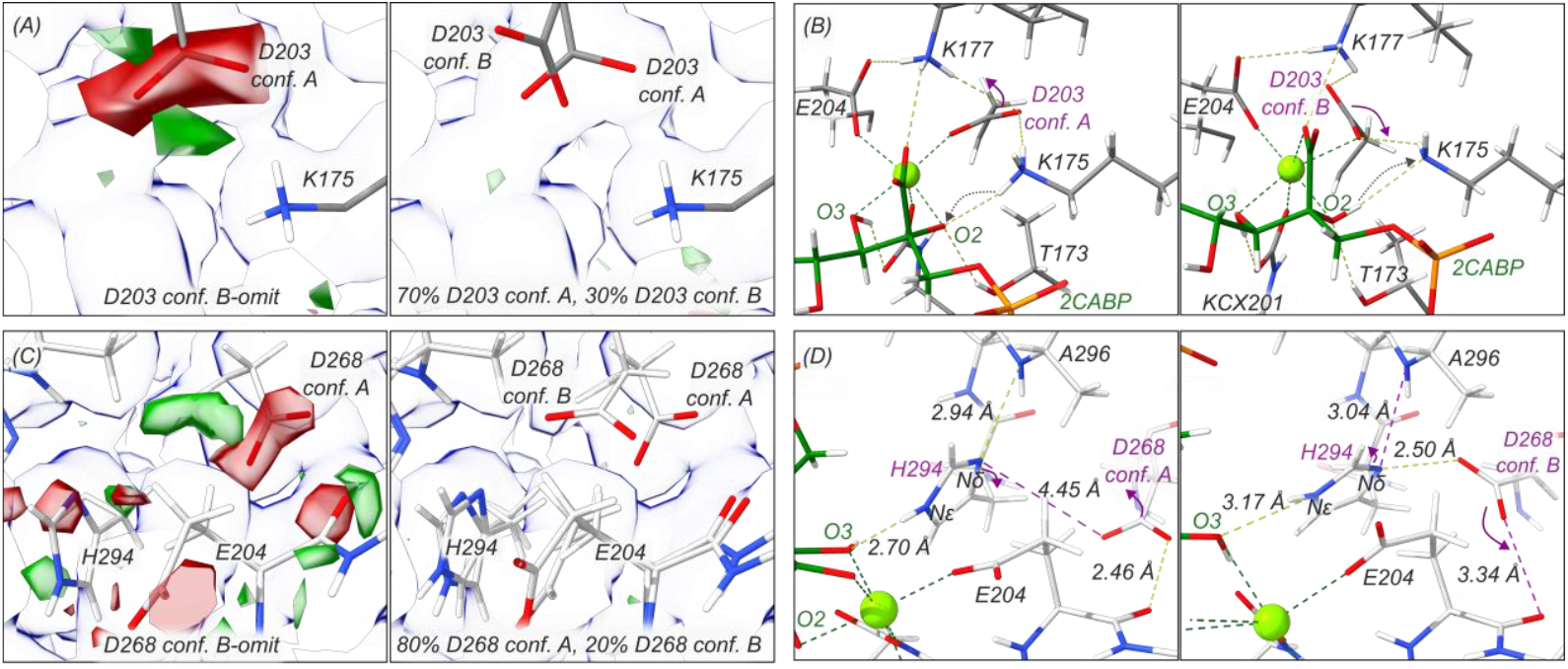
Alternative active site residue conformations linked to proton transfers. **(A)** F_o_-F_c_ omit map (left; red/green; absolute levels −1.3, 1.3) and unsharpened map (blue; absolute map level 0.01) from a “C” active site generated using a model lacking the D203 conformation B. Positive (green) densities indicate that a second D203 conformation should be modelled, while the negative (red) densities indicate that the conformation A is over-modelled. Modelling D203 in conformations A and B with 70% and 30% occupancies, respectively, removes the difference peaks (right). **(B)** Changes in the hydrogen bond network caused by the rotation of D203. Our QM/MM calculations suggest that the D203 rotation is due to proton transfer (dashed arrows) from K175 to 2CABP-O2 (see Supplementary Fig. 8). **(C)** F_o_-F_c_ omit map (left; red/green; absolute map levels −2, 2) and unsharpened map (blue; absolute map level 0.0121) from the “O” active site generated using a model lacking the D268 conformation B. Positive (green) densities indicate that a second D268 conformation should be modelled, while the negative (red) densities indicate that the positions are over-modelled. Modelling D268 in conformations A and B with 80% and 20% occupancies, respectively, removes the difference peaks (right). **(D)** Changes in hydrogen bond distances in response to the alternate conformations adopted by D268 in the “O” active site.

While the calculations support our experimentally determined protonation states, the observed protonation pattern is unexpected. The missing 2CABP-O2 proton indicates that 2CABP binding involves at least one proton transfer reaction, given that both O2 and O3 are expected to be protonated (neutral) in solution at pH 8.0 (p*K*_a_~11-12). Furthermore, our data suggest that H294 and D268 are involved in an additional proton transfer upon ligand binding. In all active sites, the major conformation of D268 places it 2.46 Å from the backbone carbonyl of E204 and 4.45 Å from H294-Nδ, forming a strong hydrogen bond with E204 but too far for hydrogen bonding with H294-Nδ (conformation A; Fig. 5C, D). In the “*C*” active sites, this is the only observed conformation of D268, but in the “*O*” active site, D268 also adopts a second conformation with the carboxylate sidechain located 2.50 Å from H294-Nδ (conformation B; ~20% occupancy; Fig. 5C, D). This distance indicates a relatively strong hydrogen bond between H294-Nδ and D268, in contrast to conformation A. The fact that we only observe conformation B of D268 in the “*O*” state suggests that H294-Nδ transfers its proton to D268 as part of the ligand binding process.

## Discussion

### Implications for the Rubisco Mechanism

The interpretation of these observations hinges on the initial proton configuration. For example, if K175 is initially unprotonated and Nδ and Nε of H294 are protonated, two proton transfers are needed to produce the observed protonation configuration (Fig. 2A, B, 6A): one from 2CABP-O2 to K175, and a second one from H294-Nδ to D268. However, we note two inconsistencies with this “K175 mechanism” (Fig. 6A). First, given that the RuBP-O2 is unprotonated at its carbonyl oxygen and that the RuBP-O2 accumulates negative charge during the enolization reaction (the first reaction step; Fig. 1E), it is likely that K175 is initially protonated, thus stabilizing the charge accumulation on O2. Second, this model suggests that the interaction between H294-Nε and 2CABP-O3 induces proton transfer from H294-Nδ to D268 (Fig. 6A), but this interaction would draw electron density away from H294, making H294-Nδ less acidic.

**Figure 6.**
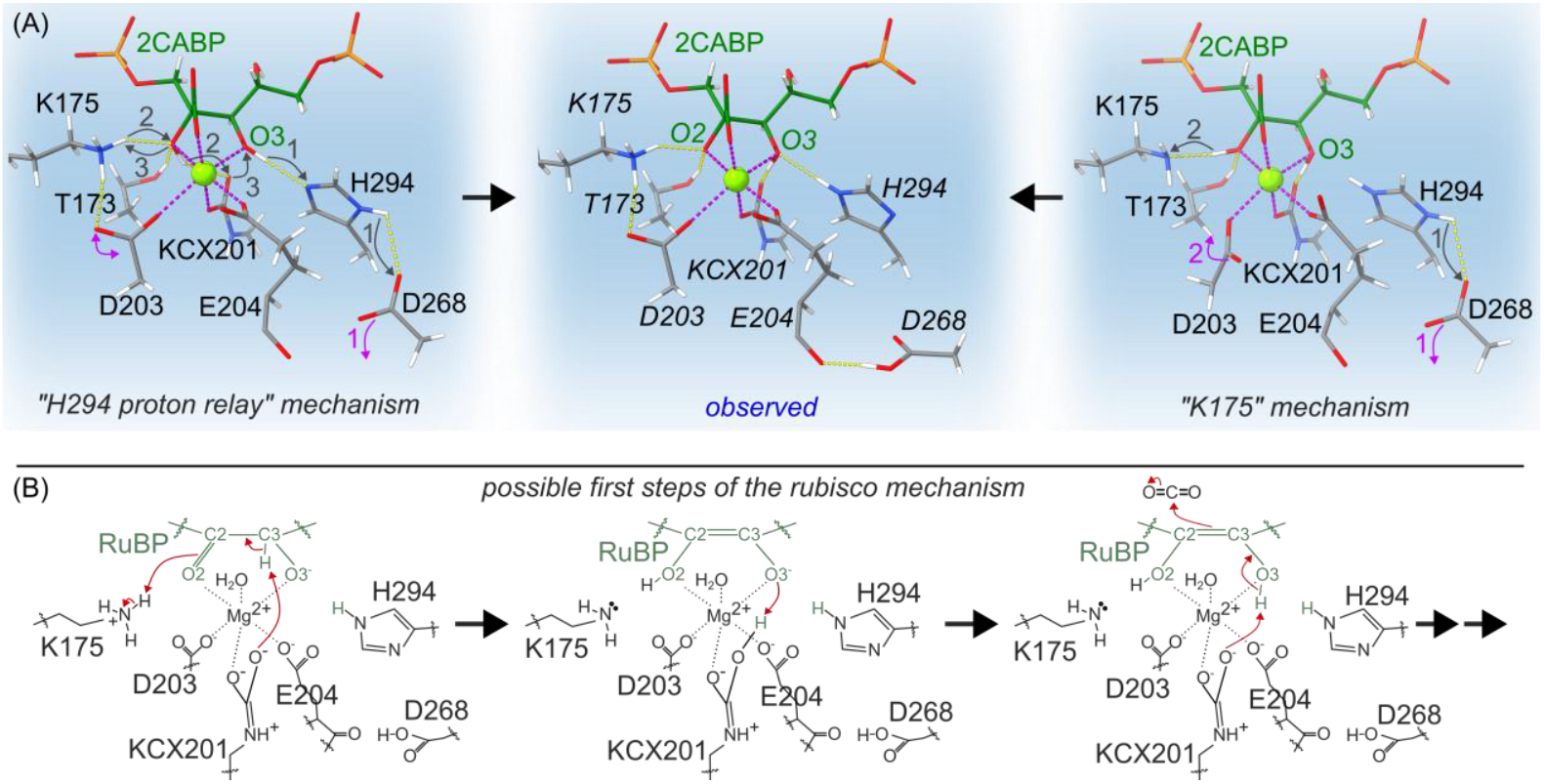
Mechanistic implications of the observed protonation configuration. **(A)** Two possible 2CABP binding mechanisms based on the observed protonation configuration. Gray arrows indicate the movement of protons. Magenta arrows indicate the movement of amino acid sidechains. **(B)** Possible first steps of the rubisco mechanism based on the observed protonation pattern of the rubisco-Mg^2+^-2CABP complex. The red arrows indicate the movement of electrons. Pink dashed lines indicate Mg^2+^ (green sphere) coordination and yellow dashed lines indicate hydrogen bonds.

We propose an alternative “H294 proton relay mechanism” in which K175 and H294-Nδ are initially protonated and H294-Nε is unprotonated (Fig. 6A). Upon 2CABP binding, H294 accepts the 2CABP-O3 proton at Nε before transferring its Nδ proton to D268, which then rotates away from H294 to form the observed hydrogen bond with the backbone carbonyl of E204. Similar histidine relays, catalysing proton exchange across the imidazole ring, have been previously reported in various systems.^27–29^ The final proton transfer step from O2 to O3 is then facilitated by KCX201, and likely accompanied by a proton exchange between K175 and 2CABP-O2 to prevent the formation of a transient doubly-unprotonated 2CABP [O2^−^, O3^−^] intermediate. Intriguingly, the X-ray structure of the spinach rubisco-RuBP complex (1RXO) shows D268 in conformation A,^9^ suggesting that the same H294 proton relay is active during RuBP binding. Under this scheme, H294 is neutral rather than positively charged, weakening the argument that RuBP-O3 is not sufficiently electronegative for a step-wise carboxylation and hydration mechanism^10^ and potentially rationalizing ^12^C/^13^C kinetic isotope effect data which show that the proton abstracted from RuBP-C3 during the enolization reaction is involved in the carboxylation transition state.^12^ KCX201 could transfer the RuBP-C3 proton to the negatively charged RuBP-O3^−^, then remove it again to facilitate the CO_2_ addition reaction (Fig. 6B). Such a “futile” proton transfer is not inconceivable given the number of protonation equilibria involved in rubisco catalysis (Supplementary Fig. 6, 8).^12,30^

## Conclusion

The atomic resolution of our cryo-EM data on the rubisco-Mg^2+^-2CABP complex has revealed surprising dynamics and an unexpected active site proton configuration, providing important insights into the energetics and structural changes induced by ligand (2CABP) binding in spinach rubisco. Our data suggests that the tight binding of 2CABP in the active site, exhibiting a femtomolar dissociation constant,^28^ does not require active site closure. Once bound, the active site samples nearly isoenergetic *open* and *closed* states, indicating that the favourable enthalpic interactions formed during the progressive ordering of the active site as it transitions from the open (*ooo*) to closed (*ccc*) state are compensated by the incurred entropy losses, which result from restraining the motions of 2CABP, loop 6, and the C-terminus. The observed structural changes appear to be connected to complex protonation equilibria and the displacement of the putative substrate water from the active site.

Similar conformational equilibria likely exist for rubisco in complex with RuBP, or CKABP (or its oxygenation reaction equivalent), yet with shifted energy levels. If so, the observations reported here imply that key assumptions made in contemporary mechanistic proposals may need to be revised to account for both the observed protonation states of active site amino acids, as well as the rich dynamics induced through a well-choreographed conformational dance between rubisco and its substrate. In this context, the perplexing contributions of distant mutations^31^ and the small subunit^3^ to rubisco’s catalytic properties may be rationalized as alterations to conformational sampling benefitting CO_2_ fixation catalysis, suggesting that rational design strategies aimed at tuning rubisco’s conformational landscapes^32^ may yield more success than traditional point mutations in the active site while complementing and rationalizing the positive results obtained via protein grafting techniques^33^ and directed evolution platforms.^34–37^ Finally, this study exemplifies how the combination of atomic resolution single particle cryo-EM and large-scale hybrid quantum/classical (QM/MM) calculations can be employed for gaining detailed insight into enzyme mechanisms.

## Methods

### Sample preparation

Market spinach (*Spinacia oleracea*) was purified, and the activated rubisco-Mg^2+^-2CABP complex was crystallized as described previously.^38,39^ One large crystal was washed three times with its mother liquor to remove any un-crystallized protein, then the crystal was dissolved in a cryo-EM buffer containing 100 mM HEPES at pH 8.0, 1.5 mM MgCl_2_, 50 mM NaHCO_3_, and 150 μM EDTA. Once dissolved, a 4mL 100 kDa molecular weight cutoff Amicon Ultra Centrifugal Filter (Millipore) was used to fully exchange any remaining crystallization buffer with the cryo-EM buffer before adjusting the rubisco concentration to 1 mg mL^−1^.

### Cryo-EM specimen preparation

HexAuFoil grids (Quantifoil) were glow-discharged in a Pelco easiGlow (25 mA, 90 s, 0.4 mbar of residual ambient air, negative polarity) before 4 uL of the 1 mg mL^−1^ rubisco-Mg^2+^-2CABP complex solution was applied on the holey gold foil side. The blotting and plunge-freezing were performed with a Vitrobot Mark IV device, with the chamber at 4ºC and 100% relative humidity, 4 s of blotting (at blot force 10) before vitrification in liquid ethane.

### Cryo-EM sample screening and data collection

Grids were examined using a Thermo Scientific Krios 5 Cryo-TEM instrument, operated at 300 kV and equipped with an E-CFEG (cold Field Emission Gun) electron source, a Selectris-X energy filter, a Falcon 4i direct electron detector, and Smart EPU software. Plasmon peak imaging mode was employed in EPU to better identify ice thickness gradients, facilitating the selection of optimal grid areas on HexAuFoil grids^40^. Following the identification of ideal imaging regions, a total of 67,056 movies were recorded in EER imaging format^41^ at a nominal magnification of 165,000x, resulting in a raw pixel size of 0.456 Å. An energy filtering slit of 10 eV was applied, with a total dose of approximately 53 e/Å^2^ administered to the sample. The nominal defocus range was set between −0.8 to −0.3 µm, in increments of 0.1 µm. Additionally, the extended aberration-free image shift (AFIS) method was utilized, allowing data collection from an entire hexagon (~20 µm cluster radius) solely by shifting the beam and by correcting for optical aberrations. The autofocus routine was set to be executed every 12 µm.

### Pre-processing, particle picking and globally averaged 3D refinement

Image processing was performed with CryoSPARC version 4.6.^42^ A summary of the processing strategy is shown in Supplementary Figure 1. Raw movies were motion corrected using the Patch Motion Correction job with default parameters. CTF parameters were determined using the Patch CTF estimation job with default parameters. 45,667 micrographs were selected from the original dataset following a certain set of quality requirements (CTF fit resolution better than 5.5 Å, relative ice thickness within 1-1.2 range, underfocus below 1.7 µm). The selected micrographs were subjected to Template picker to pick 9,719,948 particles and filtered to 6,624,751 particles based on the NCC scores. Next, the particles were extracted using 600 px box size and Fourier cropped to 100 px (4.08 Å/px), resulting in 5,136,100 particles which were submitted to reference-free 2D classification (200 classes, circular mask diameter 140 Å). 2,460,569 particles (48%) were retained based on features of their 2D class averages. These particles were submitted to another reference-free 2D classification (200 classes, circular mask diameter 140 Å), from which 2,286,371 particles (93%) were retained. These particles were then sorted into 4 classes by *ab initio* reconstruction, followed by three consecutive rounds of heterogeneous refinement using the best class from each run as input for the subsequent run. 1,632,169 particles (71%) were re-extracted in a box 600 px wide Fourier cropped to 500 px (0.548 Å/px). Extracted particles were split into Exposure Groups to cluster them based on 1-hour data collection intervals as defined in the micrograph file names. An additional explorative refinement with particles clustered to groups based on image-beam shifts did not improve results, indicating uniform optical aberration-correction over the full extended AFIS range. The particles were subjected to homogeneous refinement with D4 symmetry, optimisation of per-particle scales, optimization of per-group CTF parameters, and EWS correction with positive curvature. This refinement reached 1.33 Å resolution. A fitted Cs value of 2.67 indicated a negligible (0.26%) pixel size error. This reconstruction was used to perform reference-based motion correction^43^ (estimating hyperparameters and empirical dose weights) of the entire particle stack, from which 1,630,841 particles were subjected to a second homogeneous refinement with the same parameters as the first refinement. This refinement reached 1.25Å resolution.

### Classification of subunits with different conformations

Using the “molmap” command in ChimeraX^44^ with a resolution of 15 Å, a mask was generated around one L2S2 unit (one RbcL dimer plus two RbcS subunits, see Figure 1B). The mask was imported into CryoSPARC, binarized with a threshold of 0.08, dilated with a radius of 3 px, and soft padded with a width of 14 px. The set of 1,630,841 rubisco particles used in the globally averaged, D4-symmetric reconstruction was expanded with C4 symmetry to generate particle metadata corresponding to all 6,523,364 L2S2 sub-complexes in this set (= 4 × 1,630,841). We then ran a six-class, masked 3D classification job without alignments on the set of L2S2 particles. This approach assumes that the global alignments are valid which, given the high resolution of the D4-symmetric reconstruction, is accurate. This procedure sorted all L2S2 sub-complexes based on their conformational heterogeneity. One class showed signs of air-water interface damage and was not processed further, whereas the remaining five classes do not show any apparent damage. These five classes were subjected to signal subtraction, reconstruction, and local refinement jobs in CryoSPARC to generate five new maps for model building. Three of these maps appeared redundant, and only the highest resolution map was used for modelling this conformation (1.41 Å “C-C” model). The remaining two maps were also modelled (1.46 Å “C-P” and 1.47 Å “C-O” models), allowing the determination of three unique conformational states.

### Atomic model building and refinement

ChimeraX was used to rigid-body fit the X-ray crystal structure of activated spinach rubisco with 2CABP (PDB id 8RUC) to the unsharpened D4-symmetrized map generated in the final homogeneous refinement in CryoSPARC. Servalcat^21^ was then used to automatically refine the input model against the map, generating sharpened (F_o_) and difference (F_o_-F_c_) maps, which were used for manual model building in Coot^45^. The first Servalcat refinement was run with isotropic B-factors and riding hydrogens (Hydrogens = all), while the final Servalcat refinement was run with anisotropic B-factors and H-atom refinement (Hydrogens = yes). The final D-symmetrized model was used to create an L2S2 starting model for the three L2S2 refinements. The same model building and refinement procedure was used for the L2S2 models. Upon completion of the models, all hydrogens were stripped from the models and Servalcat was used to generate an H-omit difference map.

### Hybrid QM/MM calculations

Hybrid quantum/classical (QM/MM) calculations were performed based on the consensus cryo-EM structure of rubisco, determined at 1.25 Å resolution in the presence of the 2CABP transition-state analogue. To this end, the L1S1 unit was duplicated and aligned based on the 1.41 Å ‘C-C’ state structure for creating an L2S2 tetramer unit, with an intact protein environment around the active site. The system was solvated and neutralized with 150 mM NaCl ions. A short (1 ps) classical molecular dynamics (MD) simulation was performed at *T*=310 K, *p*=1 bar, and using a 2 fs integration timestep to relax the surrounding solvent, while keeping the protein, cofactors, and resolved water molecules fixed at their experimentally determined positions. The resulting system was trimmed to *ca*. 60,000 atoms for the subsequent QM/MM calculations. The QM region comprised K175, K177, KCX201, D203, E204, T173, H294, R295, H298, H327, K334, L335, S379, Q401 of subunit L, and E60, T65, W66 and N123 of subunit L’, 2CABP, and Mg^2+^, together with 17 water molecules, resulting in a system with 311-312 atoms (see Supplementary Table 1), with link atoms introduced between the Cα and Cβ positions of the protein residues. The QM region was modelled at the B3LYP-D3/def2-SVP(C,O,N,P,H)/def2-TZVP(Mg^2+^) level^46–49^, the MM system was described by the CHARMM36 force field^50^, with the QM/MM-coupling modelled using an additive electrostatic embedding scheme. During QM/MM optimizations, the position of heavy atoms was fixed to their experimental positions, while all hydrogen atoms within 10 Å radius around QM atoms were relaxed. Moreover, additional QM/MM models were constructed in which sidechain positions were allowed to relax during the optimization. The QM/MM optimisations were performed in multiple protonation states for comparison to the experimentally determined states (see Supplementary Table 6). The initial classical relaxation was performed using NAMD v2.14-3.0^51^, while QM/MM calculations were performed with CHARMM c38b1^52^/TURBOMOLE 7.7^53^, coupled by a Python interface^54^.

### Analysis of calculated electrostatic potential maps

Electrostatic potential (ESP) maps were calculated based on the QM/MM optimised structures at the B3LYP-D3/def2-TZVP level^46–49^ for all QM atoms and in the presence of the classical point charges. To compare with the experimental cryo-EM map, the QM/MM computed ESP maps were filtered with a Gaussian function of the form:

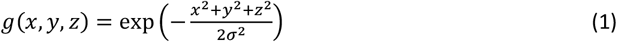

where *x,y,z* represent the electrostatic map coordinates with *σ*=0.5. In addition, difference maps between QM/MM potentials and cryo-EM maps were calculated by normalizing the root-mean-square (RMS) deviation of the subtracted maps to account for the different threshold levels. The excess difference potential around specific regions was quantified (Supplementary Fig. 7, 8) to assign the experimentally most likely protonation states. Electrostatic potential maps were calculated using TURBOMOLE 7.7^53^, while post-processing and visualization of the QM/MM-ESP and cryo-EM maps were performed using UCSF ChimeraX^44^.

## Data availability

The atomic models and cryo-EM maps have been deposited in the Protein Data Bank (PDB) and Electron Microscopy Data Bank (EMDB), respectively, with the following accession codes: 1.25 Å “consensus” structure (9TED, EMD-55825), 1.41 Å “C-C” structure (9TEE, EMD-55826), 1.46 Å “C-P” structure (9TEF, EMD-55827), and 1.47 Å “C-O” structure (9TEG, EMD-55828).

## Acknowledgements

We acknowledge the use of the Cryo-EM Uppsala facility for specimen vitrification, data storage, and computing, funded by the Department of Cell and Molecular Biology, the Disciplinary Domains of Science and Technology and of Medicine and Pharmacy at Uppsala University. We thank the National Academic Infrastructure for Supercomputing in Sweden (NAISS 2025/1-33) for computational resources. This work is supported by the Swedish Foundation for Strategic Research (SSF) via the Swedish national graduate school in neutron scattering (SwedNESS; N.C. and J.M.), Kempestiftelserna (JCSMK24-599 to JM), the Swedish Research Council (2019-03700_VR and 2023-05296_VR to C.B.), the Swedish Research Council for Sustainable Development (2019-01171_Formas to C.B.), the Knut and Alice Wallenberg Foundation (2019.0251 and 2024.0220 to V.R.I.K), the Göran Gustafsson Foundation for Research in Natural Sciences and Medicine (to V.R.I.K.), and the Swedish Research Council (2020-04081 and 2025-04607 to V.R.I.K.). I.A. is supported by the Czech Science Foundation, GACR (project No. 24-10671S to I.A.), and the Johannes Amos Comenius Operational Programme (OPJAK) financed by the European Structural and Investment Funds and the Czech Ministry of Education, Youth and Sports for the SENDISO project No. CZ.02.01.01/00/22_008/0004596. A.B. and J.K. are supported by the Department of Energy, Office of Science, Basic Energy Sciences, Division of Chemical Sciences, Geosciences, and Biosciences (DE-AC02-05CH11231).

## Author Contributions

N.C. and J.M. conceived the experiment; N.C. performed all biochemistry with initial support from I.A.; I.A. provided the 2CABP; G.G. and N.C. vitrified grids; S.M. and O.R. collected and processed the cryo-EM data with input from N.C. and G.G.; N.C. refined the models with support from G.G., A.B., and J.K.; N.C. interpreted the models; G.G. counted detected hydrogen atoms and generated OccuPy figures; P.S. and V.R.I.K. performed and interpreted the QM/MM calculations; N.C. and J.M. wrote the manuscript with input from all authors; I.A., V.R.I.K., C.B., and J.M. provided supervision and funding.

## Additional information

**Supplementary information** is available for this paper at (doi).

## References

1. Brändén, C.-I., Lindqvist, Y. & Schneider, G. Protein engineering of rubisco. Acta Crystallogr B Struct Sci 47, 824–835 (1991).

2. Parry, M. A. J. et al. Rubisco activity and regulation as targets for crop improvement. J Exp Bot 64, 717–730 (2013).

3. Mao, Y. et al. The small subunit of Rubisco and its potential as an engineering target. J Exp Bot 74, 543–561 (2022).

4. Prywes, N., Phillips, N. R., Tuck, O. T., Valentin-Alvarado, L. E. & Savage, D. F. Rubisco Function, Evolution, and Engineering. Annual Review of Biochemistry 92, 385–410 (2023).

5. Schneider, G., Lindqvist, Y. & Branden, C. I. Rubisco: Structure and Mechanism. Annual Review of Biophysics and Biomolecular Structure 21, 119–143 (1992).

6. Andersson, I. Catalysis and regulation in Rubisco. Journal of Experimental Botany 59, 1555–1568 (2007).

7. Andersson, I. & Backlund, A. Structure and function of Rubisco. Plant Physiology and Biochemistry 46, 275–291 (2008).

8. Cummins, P. L. & Gready, J. E. Kohn–Sham Density Functional Calculations Reveal Proton Wires in the Enolization and Carboxylase Reactions Catalyzed by Rubisco. J. Phys. Chem. B 124, 3015–3026 (2020).

9. Taylor, T. C. & Andersson, I. The structure of the complex between rubisco and its natural substrate ribulose 1,5-bisphosphate. Journal of Molecular Biology 265, 432–444 (1997).

10. Cleland, W. W., Andrews, T. J., Gutteridge, S., Hartman, F. C. & Lorimer, G. H. Mechanism of Rubisco: The Carbamate as General Base. Chemical Reviews 98, 549–561 (1998).

11. Tcherkez, G. G. B. et al. D2O Solvent Isotope Effects Suggest Uniform Energy Barriers in Ribulose-1,5-bisphosphate Carboxylase/Oxygenase Catalysis. Biochemistry 52, 869–877 (2013).

12. Bathellier, C. et al. Ribulose 1,5-bisphosphate carboxylase/oxygenase activates O2 by electron transfer. Proceedings of the National Academy of Sciences 117, 24234–24242 (2020).

13. Bowes, G., Ogren, W. L. & Hageman, R. H. Phosphoglycolate production catalyzed by ribulose diphosphate carboxylase. Biochemical and Biophysical Research Communications 45, 716–722 (1971).

14. Andrews, T. J., Lorimer, G. H. & Tolbert, N. E. Ribulose diphosphate oxygenase. I. Synthesis of phosphoglycolate by fraction-1 protein of leaves. Biochemistry 12, 11–18 (1973).

15. Lorimer, G. H. & Miziorko, H. M. Carbamate formation on the.epsilon.-amino group of a lysyl residue as the basis for the activation of ribulosebisphosphate carboxylase by carbon dioxide and magnesium(2+). Biochemistry 19, 5321–5328 (1980).

16. Knight, S., Andersson, I. & Brindn, C.-I. Crystallographic Analysis of Ribulose 1,5-Bisphosphate Carboxylase from Spinach at 2.4 A Resolution Subunit Interactions and Active Site. Journal of Molecular Biology 215, 113–160 (1990).

17. Andersson, I. Large Structures at High Resolution: The 1.6 Å Crystal Structure of Spinach Ribulose-1,5-Bisphosphate Carboxylase/Oxygenase Complexed with 2-Carboxyarabinitol Bisphosphate. Journal of Molecular Biology 259, 160–174 (1996).

18. Blakeley, M. P. Neutron macromolecular crystallography. Crystallography Reviews 15, 157–218 (2009).

19. Nakane, T. et al. Single-particle cryo-EM at atomic resolution. Nature 587, 152–156 (2020).

20. Hussein, R. et al. Cryo–electron microscopy reveals hydrogen positions and water networks in photosystem II. Science 384, 1349–1355 (2024).

21. Yamashita, K., Palmer, C. M., Burnley, T. & Murshudov, G. N. Cryo-EM single-particle structure refinement and map calculation using Servalcat. Acta Cryst D 77, 1282–1291 (2021).

22. Yip, K. M., Fischer, N., Paknia, E., Chari, A. & Stark, H. Atomic-resolution protein structure determination by cryo-EM. Nature 587, 157–161 (2020).

23. Schloss, J. V. Comparative affinities of the epimeric reaction-intermediate analogs 2- and 4-carboxy-D-arabinitol 1,5-bisphosphate for spinach ribulose 1,5-bisphosphate carboxylase. Journal of Biological Chemistry 263, 4145–4150 (1988).

24. Duff, A. P. et al. The Transition between the Open and Closed States of Rubisco is Triggered by the Inter-Phosphate Distance of the Bound Bisphosphate. Journal of Molecular Biology 298, 903–916 (2000).

25. Taylor, T. C. & Andersson, I. Structural transitions during activation and ligand binding in hexadecameric Rubisco inferred from the crystal structure of the activated unliganded spinach enzyme. Nat Struct Mol Biol 3, 95–101 (1996).

26. Pierce, J., Tolbert, N. E. & Barker, R. Interaction of ribulosebisphosphate carboxylase/oxygenase with transition-state analogs. Biochemistry 19, 934–942 (1980).

27. Azadmanesh, J., Lutz, W. E., Coates, L., Weiss, K. L. & Borgstahl, G. E. O. Direct detection of coupled proton and electron transfers in human manganese superoxide dismutase. Nat Commun 12, 2079 (2021).

28. Bhowmick, A. et al. Structural evidence for intermediates during O2 formation in photosystem II. Nature 617, 629–636 (2023).

29. Mühlbauer, M. E. et al. Water-Gated Proton Transfer Dynamics in Respiratory Complex I. J. Am. Chem. Soc. 142, 13718–13728 (2020).

30. Bathellier, C. & Tcherkez, G. Experimental evidence for extra proton exchange in ribulose 1,5-bisphosphate carboxylase/oxygenase catalysis. Communicative & Integrative Biology 15, 68–74 (2022).

31. Whitney, S. M. et al. Isoleucine 309 acts as a C4 catalytic switch that increases ribulose-1,5-bisphosphate carboxylase/oxygenase (rubisco) carboxylation rate in Flaveria. Proceedings of the National Academy of Sciences 108, 14688–14693 (2011).

32. Damry, A. M. & Jackson, C. J. The evolution and engineering of enzyme activity through tuning conformational landscapes. Protein Eng Des Sel 34, gzab009 (2021).

33. Zhou, Y., Gunn, L. H., Birch, R., Andersson, I. & Whitney, S. M. Grafting Rhodobacter sphaeroides with red algae Rubisco to accelerate catalysis and plant growth. Nat. Plants 9, 978–986 (2023).

34. Gionfriddo, M. et al. Laboratory evolution of Rubisco solubility and catalytic switches to enhance plant productivity. Nat. Plants 1–12 (2025).

35. Hoffmann, U. A. et al. A cyanobacterial screening platform for Rubisco mutant variants. Preprint at 10.1101/2025.01.22.633911 (2025).

36. McDonald, J. L., Shapiro, N. P., Whitney, S. M. & Wilson, R. H. In vivo directed evolution of an ultra-fast RuBisCO from a semi-anaerobic environment imparts oxygen resistance. (2025).

37. Prywes, N. et al. A map of the rubisco biochemical landscape. Nature 1–6 (2025) doi:10.1038/s41586-024-08455-0.

38. Andersson, I., Tjäder, A. C., Cedergren-Zeppezauer, E. & Brändén, C. I. Crystallization and preliminary x-ray studies of spinach ribulose 1,5-bisphosphate carboxylase/oxygenase complexed with activator and a transition state analogue. Journal of Biological Chemistry 258, 14088–14090 (1983).

39. Andersson, I. & Brändén, C.-I. Large single crystals of spinach 1,5-bisphosphate carboxylase/oxygenase suitable for X-ray studies. Journal of Molecular Biology 172, 363–366 (1984).

40. Hagen, W. J. H. Light ‘Em up: Efficient Screening of Gold Foil Grids in Cryo-EM. Front Mol Biosci 9, 912363 (2022).

41. Guo, H. et al. Electron-event representation data enable efficient cryoEM file storage with full preservation of spatial and temporal resolution. IUCrJ 7, 860–869 (2020).

42. Punjani, A., Rubinstein, J. L., Fleet, D. J. & Brubaker, M. A. cryoSPARC: algorithms for rapid unsupervised cryo-EM structure determination. Nat Methods 14, 290–296 (2017).

43. Zivanov, J., Nakane, T. & Scheres, S. H. W. A Bayesian approach to beam-induced motion correction in cryo-EM single-particle analysis. IUCrJ 6, 5–17 (2019).

44. Pettersen, E. F. et al. UCSF ChimeraX: Structure visualization for researchers, educators, and developers. Protein Science 30, 70–82 (2021).

45. Emsley, P., Lohkamp, B., Scott, W. G. & Cowtan, K. Features and development of Coot. Acta Crystallogr D Biol Crystallogr 66, 486–501 (2010).

46. Becke, A. D. Density-functional thermochemistry. III. The role of exact exchange. The Journal of Chemical Physics 98, 5648–5652 (1993).

47. Lee, C., Yang, W. & Parr, R. G. Development of the Colle-Salvetti correlation-energy formula into a functional of the electron density. Phys. Rev. B 37, 785–789 (1988).

48. Grimme, S., Antony, J., Ehrlich, S. & Krieg, H. A consistent and accurate ab initio parametrization of density functional dispersion correction (DFT-D) for the 94 elements H-Pu. J. Chem. Phys. 132, 154104 (2010).

49. Schäfer, A., Horn, H. & Ahlrichs, R. Fully optimized contracted Gaussian basis sets for atoms Li to Kr. The Journal of Chemical Physics 97, 2571–2577 (1992).

50. Best, R. et al. Optimization of the Additive CHARMM All-Atom Protein Force Field Targeting Improved Sampling of the Backbone phi, psi and Side-Chain chi(1) and chi(2) Dihedral Angles. J. Chem. Theory Comput. 8, 3257–3273 (2012).

51. Phillips, J. C. et al. Scalable molecular dynamics on CPU and GPU architectures with NAMD. J. Chem. Phys. 153, 044130 (2020).

52. Brooks, B. et al. CHARMM: The Biomolecular Simulation Program. Journal of Computational Chemistry 30, 1545–1614 (2009).

53. Balasubramani, S. G. et al. TURBOMOLE: Modular program suite for ab initio quantum-chemical and condensed-matter simulations. J. Chem. Phys. 152, 184107 (2020).

54. Riahi, S. & Rowley, C. N. The CHARMM-TURBOMOLE interface for efficient and accurate QM/MM molecular dynamics, free energies, and excited state properties. J. Comput. Chem. 35, 2076–2086 (2014).

