## Supplementary-Information for "Atomic resolution structure of spinach rubisco reveals protons and dynamics"

***Supplementary Discussion***

*(Page 2) S1. Counting detected hydrogen atoms*
*(Page 3) S2. Estimating occupancy and number of active sites in each active site state*
*(Page 5) S3. 2CABP binding models*
*(Page 6) S4. Source of the conformational heterogeneity*

***Supplementary Tables***

*(Page 7) Supplementary Table 1. Counting detected H-atoms*
*(Page 8) Supplementary Table 2. Summary of 6 classes obtained from 3D classification.*
*(Page 8) Supplementary Table 3. Estimated occupancies of structural components involved in active site closure.*
*(Page 9) Supplementary Table 4. Estimated # active sites in each active site state.*
*(Page 10) Supplementary Table 5. 2CABP binding equilibria.*
*(Page 11) Supplementary Table 6. List of explored protonation states in QM/MM calculations.*
*(Page 12) Supplementary Table 7. Geometries of first-coordination sphere from optimised QM/MM calculations.*

***Supplementary Figures***

*(Page 13) Supplementary Fig. 1. Atomic resolution data.*
*(Page 14) Supplementary Fig. 2. Cryo-EM data processing overview.*
*(Page 15) Supplementary Fig. 3. Air-water interface damage.*
*(Page 16) Supplementary Fig. 4. Occupancy estimation method.*
*(Page 17) Supplementary Fig. 5. QM/MM setup and comparison of cryo-EM and QM/MM ESP maps.*
*(Page 18) Supplementary Fig. 6. QM/MM optimised geometries.*
*(Page 19) Supplementary Fig. 7. Comparison of cryo-EM and QM/MM difference maps.*
*(Page 20) Supplementary Fig. 8. Calculated ESP maps from multiple protonation states.*
*(Page 21) Supplementary Fig. 9. OccuPy figures.*

**Supplementary Information**

*S1. Counting detected hydrogen atoms*

Counting detected hydrogen atoms was performed as follows. Using the fully refined atomic model and experimental half-maps, an $F_o$-$F_c$ H-omit difference map was calculated with Servalcat[1] (command 'fofc'). Difference map values at hydrogen atom coordinates were then measured with ChimeraX[2] (commands 'measure mapvalue #1 atoms H attribute fofc_map_value' and 'save hydrogens_fofc-map-values.defattr attrName a:fofc_map_value'). Measured difference map values were also extracted separately for polar and non-polar hydrogen atoms (by using 'select HC' and 'select H & ~HC' in ChimeraX, respectively, before separating .defattr file with the option 'selectedOnly true'). For each $F_o$-$F_c$ H-omit difference map, absolute levels corresponding to 2 and 3 sigmas were determined with ChimeraX using the command 'volume #1 rmsLevel 2' and 'volume #1 rmsLevel 3', respectively, and reading the absolute level from the Volume Viewer panel. Measured difference map values at H atom coordinates equal to or greater than these thresholds were then counted, and divided by the number of H atoms in the corresponding atomic model to obtain the fraction of H atoms detected at each threshold.

For comparison, we also applied the same procedure to the atomic-resolution cryo-EM structures of apoferritin from Nakane et al. 2020[3] (PDB 7A4M, EMD-11638) and Yip et al 2020[4] (PDB 6Z6U, EMD-11103). All atomic models spanning only an asymmetric unit were expanded to span their entire experimental map prior to calculating the $F_o$-$F_c$ H-omit difference map.

A slightly different procedure[1] uses the PEAKMAX program from the CCP4 suite[5] to find peaks without considering H atom coordinates. The obtained peak list is then filtered to only retain those peaks with a centroid within 0.3 Å of an H atom coordinate. Our approach is functionally equivalent to the PEAKMAX approach with a distance tolerance of 0 Å. As a result, our procedure might: 1) underestimate the number of peaks, or 2) underestimate the true height of peaks for which the centroid does not exactly lie on an H atom coordinate. This makes our procedure overall slightly more conservative than the previously published one.

*S2. Estimating occupancy and number of active sites in each active site state*

To identify any conformational heterogeneity, we used C4 symmetry expansion to create C4-particles containing one L2S2 unit with two complete active sites (Fig. 1B), and performed 3D classification (without alignment) on the new C4-particles (Supplementary Fig. 1). 16% of particles were sorted into a class with slight air-water interface damage and were not processed further (Supplementary Fig. 3, Supplementary Table 2). 53% of the C4-particles were sorted into three classes with fully closed active sites and resolutions of 1.41 Å, 1.45 Å, and 1.46 Å. Using the 1.41 Å dataset, we refined one model to represent this state, hereon referred to as the *C-C* model (active site 1 - active site 2 = closed-closed). The remaining 31% of the C4-particles were sorted into two classes with mixed active site states, and we refined models for both of these classes. The reconstruction of one of these classes reaches 1.46 Å, and the model refined to this data has one closed active site and one active site with mixed N-domain and loop 6 states (the *C-P* model; *P* corresponds to the mixed N-domain state). The reconstruction of the second of the mixed active site classes reaches 1.47 Å, and the model refined to this data has one closed active site and one active site with the N-domain open and loop 6 mixed (the *C-O* model; *O* denotes the open N-domain state). We observe a weak density for some residues in the C-terminus in the closed state in the "*P*" and "*O*" active sites in the *C-P* and *C-O* models, respectively.

To estimate the occupancy of the N-domain, loop 6, and the C-terminus in each active site, the occupancies of residues 48-67 (N-domain), 330-338 (loop 6), and 463-475 (C-terminus) in their closed positions was set to 20%, 50%, 60%, 70%, 80%, 90%, and 100%, and $F_o$-$F_c$ maps were generated using the Servalcat "fofc" command.[1] The $F_o$-$F_c$ maps were then inspected at a contour level that removed most noise, and the occupancy which minimized difference peaks was chosen (or if two sequential 10% increments under- and over-estimate the occupancy, respectively, a middle value was chosen; see Supplementary Fig. 4). Supplementary Table 3 summarizes the occupancy assignments for the three L2S2 models, as well as the consensus model.

The occupancy estimations indicate that all active sites still reflect multiple states and, based on the fact that the C-terminus cannot close while loop 6 is open, we identify six possible combinations of the N-domain, loop 6, and the C-terminus: *ooo*, *coo*, *cco*, *ccc*, *oco*, and *occ*. The occupancies were then used to distribute the active sites in each model between these states. In some cases, multiple interpretations could not be distinguished. For example, the *P* active site has ~50% occupancy for both the N-domain and loop 6 in the closed position, and 20% for the C-terminus). Because the 20% with the C-terminus closed must have loop 6 closed, these active sites belong to either the *occ* or *ccc* states. We cannot determine to which state they belong, so they were split evenly between the two. For the 80% remaining active sites, one extreme interpretation is that 40% of the particles have a closed N-domain and a closed loop 6 (*cco*), and 40% have an open N-domain and an open loop 6 (*ooo*). The opposite extreme interpretation is that 40% of the particles have a closed N-domain and an open loop 6 (*coo*), and 40% have an open N-domain and a closed loop 6 (*oco*). Again, we cannot determine to which state they belong, so the active sites were split evenly among these four states.

To determine the occupancy of the full dataset, the same occupancy analysis was applied to the consensus model. From this, the occupancy of the C-terminus is estimated to be 85%, while the occupancy of loop 6 and the N-domain is estimated to be ~90%. Because the C-terminus can only be closed if loop 6 is closed, the *occ* and *ccc* states must comprise 85% of the total particles. The absolute number of *occ* states identified in the L2S2 occupancy analysis was taken to be the absolute number in the total population. This corresponds to 2% occ in the total population, leaving 83% in the *ccc* state. Similarly, the absolute number of *oco*, *ooo*, and *coo* states identified in the L2S2 occupancy analysis were also treated as the absolute number in each of these states in the total population, leaving 4% which can then be assigned to the *cco* state. This allocation of states gives 85% with the C-terminus closed, 89% with the N-domain closed, and 93% with loop 6 closed. These estimates are likely accurate within a few percentage points, which is unlikely to change the qualitative analysis (see next section) as a 2% error corresponds to ~0.6 kcal $mol^{-1}$ at 298K.

*S3. 2CABP binding models*

The equilibrium constants separating each of the active site states and the free energy differences between them were calculated using the active site estimates from the consensus structure. Of particular interest is the number of *ooo* active sites. 2CABP binding has been reported to be biphasic, whereby in the first phase, a reversible complex (EI) is quickly formed with a measured inhibition constant, $K_i$, of 0.4 x $10^{-6}$ M.[6] This is comparable to the binding of other diphosphate ligands and the Michaelis constant for the substrate, RuBP ($K_m$ = 20 µM).[7] In the second phase, slow structural rearrangements are proposed to form an "irreversible" complex (EI*) with a measured dissociation constant, $K_D$ = ~190 fM.[8] This second phase was thought to involve, minimally, the formation of hydrogen bonds between the loop 6 K334 and (1) the 2CABP carboxylate group and (2) the 2CABP P2 which "lock" the rubisco-$Mg^{2+}$-2CABP complex in the fully closed active site state. If the transition between EI and EI* is due to active site closure, the number of particles found in each active site state should be consistent with the equilibrium constant $K_2 = K_D/K_i$ = ~4 x $10^{-7}$; that is, for the ~13 million active sites in our final 1.25 Å reconstruction, only ~5 open active sites are expected (# active sites/(1 + $K_2$)). Therefore, the number of *ooo* active sites is ~$10^5$x greater than expected, which suggests that *ooo* is not EI. Supplementary Table 5 summarizes the previously reported and newly calculated equilibrium constants for the following equilibrium model:

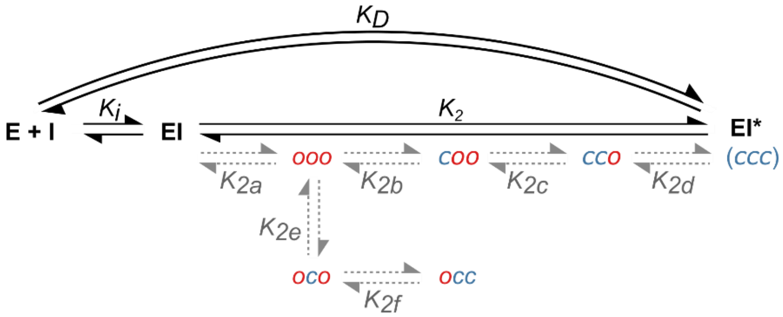

Here, we include two "off-axis" intermediates, *oco* and *occ*, formed when loop 6 closes before the N-domain, precluding the N-domain from closing. These are likely "non-physiological" states formed here because the 2CABP carboxyl group interacts with the loop 6 K334.

*S4. Source of the conformational heterogeneity*

To rule out the possibility that the open active sites were preferentially selected during the data processing, we performed a hypothetical calculation using all picked particles as our total population. In this case, the number of expected open active sites increases from 5 to 8.

To rule out partial occupancy of 2CABP as the source of the heterogeneity, we used OccuPy[9] to estimate the local scale of our cryo-EM maps in each active site (Supplementary Fig. 9). In the partially open and open active sites, the middle of the 2CABP (between the phosphates) has decreased density, but the 2CABP phosphates are at the same scale as nearby residues. We interpret this to mean that the 2CABP is fully bound, but mobile in these active sites. For this reason, we conclude that the observed heterogeneity is not the result of 2CABP release, but rather conformational sampling with 2CABP bound.

### Supplementary Table 1. Counting detected H-atoms

*The number of hydrogen atoms detected at two and three sigma in the four models obtained in this study and in the two previously published atomic resolution apoferritin models.*

| Model | Type of H | H atoms in model | Absolute level for 2 sigma | Peaks at H coordinates >=2 sigma | Fraction of H detected at 2 sigma (%) | Absolute level for 3 sigma | Peaks at H coordinates >=3 sigma | Fraction of H detected at 3 sigma (%) |
|---|---|---|---|---|---|---|---|---|
| ApoF Nakane2020 expanded | all | 32880 | 1.084 | 23352 | 71.0 | 1.626 | 19209 | 58.4 |
| | aliphatic | 24936 | 1.084 | 18240 | 73.1 | 1.626 | 14703 | 59.0 |
| | polar | 7944 | 1.084 | 5112 | 64.4 | 1.626 | 4506 | 56.7 |
| ApoF Yip2020 | all | 32784 | 0.397 | 27800 | 84.8 | 0.595 | 26887 | 82.0 |
| | aliphatic | 24840 | 0.397 | 21899 | 88.2 | 0.595 | 21214 | 85.4 |
| | polar | 7944 | 0.397 | 5901 | 74.3 | 0.595 | 5673 | 71.4 |
| Consensus - expanded | all | 36432 | 0.521 | 32419 | 89.0 | 0.781 | 31098 | 85.4 |
| | aliphatic | 28184 | 0.521 | 25235 | 89.5 | 0.781 | 24066 | 85.4 |
| | polar | 8248 | 0.521 | 7184 | 87.1 | 0.781 | 7032 | 85.3 |
| C-P | all | 9112 | 0.301 | 7658 | 84.0 | 0.451 | 7224 | 79.3 |
| | aliphatic | 7036 | 0.301 | 5913 | 84.0 | 0.451 | 5538 | 78.7 |
| | polar | 2076 | 0.301 | 1745 | 84.1 | 0.451 | 1686 | 81.2 |
| C-O | all | 8939 | 0.301 | 7222 | 80.8 | 0.451 | 6722 | 75.2 |
| | aliphatic | 6898 | 0.301 | 5561 | 80.6 | 0.451 | 5132 | 74.4 |
| | polar | 2041 | 0.301 | 1661 | 81.4 | 0.451 | 1590 | 77.9 |
| C-C | all | 9107 | 0.300 | 8077 | 88.7 | 0.451 | 7733 | 84.9 |
| | aliphatic | 7046 | 0.300 | 6274 | 89.0 | 0.451 | 5968 | 84.7 |
| | polar | 2061 | 0.300 | 1803 | 87.5 | 0.451 | 1765 | 85.6 |

### Supplementary Table 2. Summary of 6 classes obtained from 3D classification.

*The number of particles in each of the six classes obtained by 3D classification using L2S2 sub-particles. For classes 0-4, models with resolutions ranging from 1.41-1.47 Å were obtained. In each of these classes, the N-domain, loop 6, and C-terminus were assigned as closed, mixed, or open using the unsharpened maps. Classes 1, 2, and 4 were used to construct three models which, based on the N-domain state in each active site, were designated "C-P", "C-O", and "C-C".*

| Class | Particles | | Resolution | Active Sites | | N-Domain | Loop 6 | C-terminus | Model |
|---|---|---|---|---|---|---|---|---|---|
| | # | % | | Chain ID | State | | | | |
| 0 | 1.0E+06 | 16% | 1.46 Å | - | - | closed | closed | closed | - |
| | | | | - | - | closed | closed | closed | |
| **1** | **9.8E+05** | **15%** | **1.46 Å** | **A** | ***"P"*** | **mixed** | **mixed** | **open** | ***C-P*** |
| | | | | **O** | ***"C"*** | **closed** | **closed** | **closed** | |
| **2** | **1.1E+06** | **16%** | **1.47 Å** | **A** | ***"C"*** | **closed** | **closed** | **closed** | ***C-O*** |
| | | | | **O** | ***"O"*** | **open** | **mixed** | **open** | |
| 3 | 9.3E+05 | 14% | 1.45 Å | - | - | closed | closed | closed | - |
| | | | | - | - | closed | closed | closed | |
| **4** | **1.5E+06** | **23%** | **1.41 Å** | **A** | ***"C"*** | **closed** | **closed** | **closed** | ***C-C*** |
| | | | | **O** | ***"C"*** | **closed** | **closed** | **closed** | |
| 5 | 1.0E+06 | 16% | - | - | - | - | - | - | - |

### Supplementary Table 3. Estimated occupancies of structural components involved in active site closure.

*The occupancy of the N-domain, Loop 6, and C-terminus in their closed positions was estimated in the consensus model and three L2S2 models obtained by 3D classification.*

| Model | Active Sites | | N-Domain | Loop 6 | C-terminus |
|---|---|---|---|---|---|
| | Chain ID | State | | | |
| *C-C* | A | *"C"* | 100% | 100% | 90% |
| | O | *"C"* | 100% | 100% | 90% |
| *C-P* | A | *"P"* | 50% | 50% | 20% |
| | O | *"C"* | 100% | 100% | 90% |
| *C-O* | A | *"C"* | 100% | 100% | 80% |
| | O | *"O"* | 0% | 50% | 20% |
| consensus | A | - | 90% | 90% | 85% |

### Supplementary Table 4. Estimated # active sites in each active site state.

*Based on the occupancies of the N-domain, Loop 6, and the C-terminus, each active site in each model was assigned to multiple 3-letter active site states.*

| Model | Active Site | | | Occ. | N-Domain | Loop 6 | C-terminus | # Active Sites |
|---|---|---|---|---|---|---|---|---|
| | Chain ID | State | 3-letter state | | | | | |
| *C-C* | A | *"C"* | *ccc* | 90% | closed | closed | closed | 1.3E+06 |
| | | | *cco* | 10% | closed | closed | open | 1.5E+05 |
| | O | *"C"* | *ccc* | 90% | closed | closed | closed | 1.3E+06 |
| | | | *cco* | 10% | closed | closed | open | 1.5E+05 |
| *C-P* | A | *"P"* | *ccc* | 10% | closed | closed | closed | 9.8E+04 |
| | | | *cco* | 20% | closed | closed | open | 2.0E+05 |
| | | | *coo* | 20% | closed | open | open | 2.0E+05 |
| | | | *ooo* | 20% | open | open | open | 2.0E+05 |
| | | | *oco* | 20% | open | closed | open | 2.0E+05 |
| | | | *occ* | 10% | open | closed | closed | 9.8E+04 |
| | O | *"C"* | *ccc* | 90% | closed | closed | closed | 8.8E+05 |
| | | | *cco* | 10% | closed | closed | open | 9.8E+04 |
| *C-O* | A | *"C"* | *ccc* | 80% | closed | closed | closed | 8.5E+05 |
| | | | *cco* | 20% | closed | closed | open | 2.1E+05 |
| | O | *"O"* | *ooo* | 50% | open | open | open | 5.3E+05 |
| | | | *oco* | 30% | open | closed | open | 3.2E+05 |
| | | | *occ* | 20% | open | closed | closed | 2.1E+05 |
| consensus | A | - | *ccc* | 83% | closed | closed | closed | 1.1E+07 |
| | | | *cco* | 4% | closed | closed | open | 5.2E+05 |
| | | | *coo* | 2% | closed | open | open | 2.0E+05 |
| | | | *ooo* | 6% | open | open | open | 7.3E+05 |
| | | | *oco* | 4% | open | closed | open | 5.1E+05 |
| | | | *occ* | 2% | open | closed | closed | 3.1E+05 |

### Supplementary Table 5. 2CABP binding equilibria.

*The equilibrium constants associated with 2CABP binding were calculated based on the number of active sites in each state and literature values. The apparent equilibrium constants, $K_2^{app}$ and $K_D^{app}$, assume that the ooo state is EI.*

| | Value | $\Delta G$ (kcal mol$^{-1}$) | Source |
|---|---|---|---|
| $K_i$ | 4.70E-07 | 8.63 | ref. 22 |
| $K_D$ | 1.90E-13 | 17.34 | ref. 17 |
| $K_2$ | 4.04E-07 | 8.71 | $K_D$ / $K_i$ |
| $K_{2a}$ | 6.02E-06 | 7.12 | $K_2$ / ($K_{2b}$*$K_{2c}$ *$K_{2d}$) |
| $K_{2b}$ | 3.70E+00 | -0.77 | (# *ooo*) / (# *coo*) |
| $K_{2c}$ | 3.77E-01 | 0.58 | (# *coo*) / (# cco) |
| $K_{2d}$ | 4.82E-02 | 1.80 | (# *cco*) / (# *ccc*) |
| $K_{2e}$ | 1.41E+00 | -0.20 | (# *ooo*) / (# *oco*) |
| $K_{2f}$ | 1.66E+00 | -0.30 | (# *oco*) / (# *occ*) |
| $K_2^{app}$ | 6.72E-02 | 1.60 | $K_{2b}$*$K_{2c}$*$K_{2d}$ |
| $K_D^{app}$ | 3.16E-08 | 10.22 | $K_i$*$K_{2app}$ |

### Supplementary Table 6. List of explored protonation states in QM/MM calculations.

| Model | Protonation states | Optimised coordinates | Comments |
|---|---|---|---|
| Model 1 | 2CABP(O2=$O^-$/O3=O-H), $K175^+$, $K177^+$, $KCX201^-$, $D203^-$, $E204^-$, H294-ε, H298-p, H327-σ, $K334^+$, $E60^-$ | Hydrogens | Optimised state as starting model. Same protonation states as assigned experimentally |
| Model 2 | 2CABP(O2=$O^-$/O3=O-H), $K175^0$, $K177^+$, $KCX201^-$, $D203^0$, $E204^-$, H294-σ, H298-ε, H327-p, $K334^+$, $E60^-$ | Hydrogens | Upon optimisation, $K175^0$/$D203^0$ reverts to $K175^+$/$D203^-$ |
| Model 3 | 2CABP(O2=$O^-$/O3=O-H), $K175^0$, $K177^+$, $KCX201^-$, $D203^0$, $E204^-$, H294-p, H298-σ, H327- ε, $K334^+$, $E60^-$ | Hydrogens | Optimised state as starting model |
| Model 4 | 2CABP(O2=$O^-$/O3=O-H), $K175^+$, $K177^+$, $KCX201^-$, $D203^-$, $E204^-$, H294-ε, H298-p, H327-σ, $K334^+$, $E60^-$ | Hydrogens + 2CABP, KCX201, D203, E204, K175 | Optimised state as starting model. Same protonation states as assigned experimentally. |
| Model 5 | 2CABP(O2=O-H/O3=$O^-$), $K175^+$, $K177^+$, $KCX201^-$, $D203^-$, $E204^-$, H294-ε, H298-p, H327-σ, $K334^+$, $E60^-$ | Hydrogens + 2CABP, KCX201, D203, E204, K175 | Alternative protonation state of 2CABP |
| Model 6 | 2CABP(O2=O-H/O3=O-H), $K175^0$, $K177^+$, $KCX201^-$, $D203^-$, $E204^-$, H294-ε, H298-p, H327-σ, $K334^+$, $E60^-$ | Hydrogens + 2CABP, KCX201, D203, E204, K175 | Deprotonated K175 by O2 of 2CABP |
| Model 7 | 2CABP(O2=O-H/O3=O-H), $K175^+$, $K177^+$, $KCX201^-$, $D203^-$, $E204^-$, H294-ε, H298-p, H327-σ, $K334^+$, $E60^-$ | Hydrogens + 2CABP, KCX201, D203, E204, K175 | Fully protonated 2CABP and K175 |

### Supplementary Table 7. Geometries of first-coordination sphere from optimised QM/MM calculations.

*Distances are shown in units of ångströms. Model 1 corresponds to heavy atom coordinates as assigned experimentally. RMSD of the metal-ligand distances are calculated with respect the experimental coordinates.*

| Bond | Model 1 | Model 4 | Model 5 | Model 6 | Model 7 |
|---|---|---|---|---|---|
| 2CABP(O2)-$Mg^{2+}$ | 2.15 | 2.06 | 2.17 | 2.19 | 2.23 |
| 2CABP(O3)-$Mg^{2+}$ | 2.19 | 2.22 | 2.05 | 2.14 | 2.23 |
| 2CABP($COO^-$)-$Mg^{2+}$ | 1.99 | 2.09 | 2.13 | 2.10 | 2.05 |
| KCX201($COO^-$)-$Mg^{2+}$ | 2.02 | 2.05 | 2.07 | 2.07 | 2.01 |
| D203($COO^-$)-$Mg^{2+}$ | 1.97 | 2.01 | 2.05 | 1.98 | 1.97 |
| E204($COO^-$)-$Mg^{2+}$ | 2.04 | 2.10 | 1.98 | 2.06 | 2.00 |
| $d$($Mg^{2+}$-ligand) RMSD | - | 0.0251 | 0.0521 | 0.0192 | 0.0133 |

### Extended Data

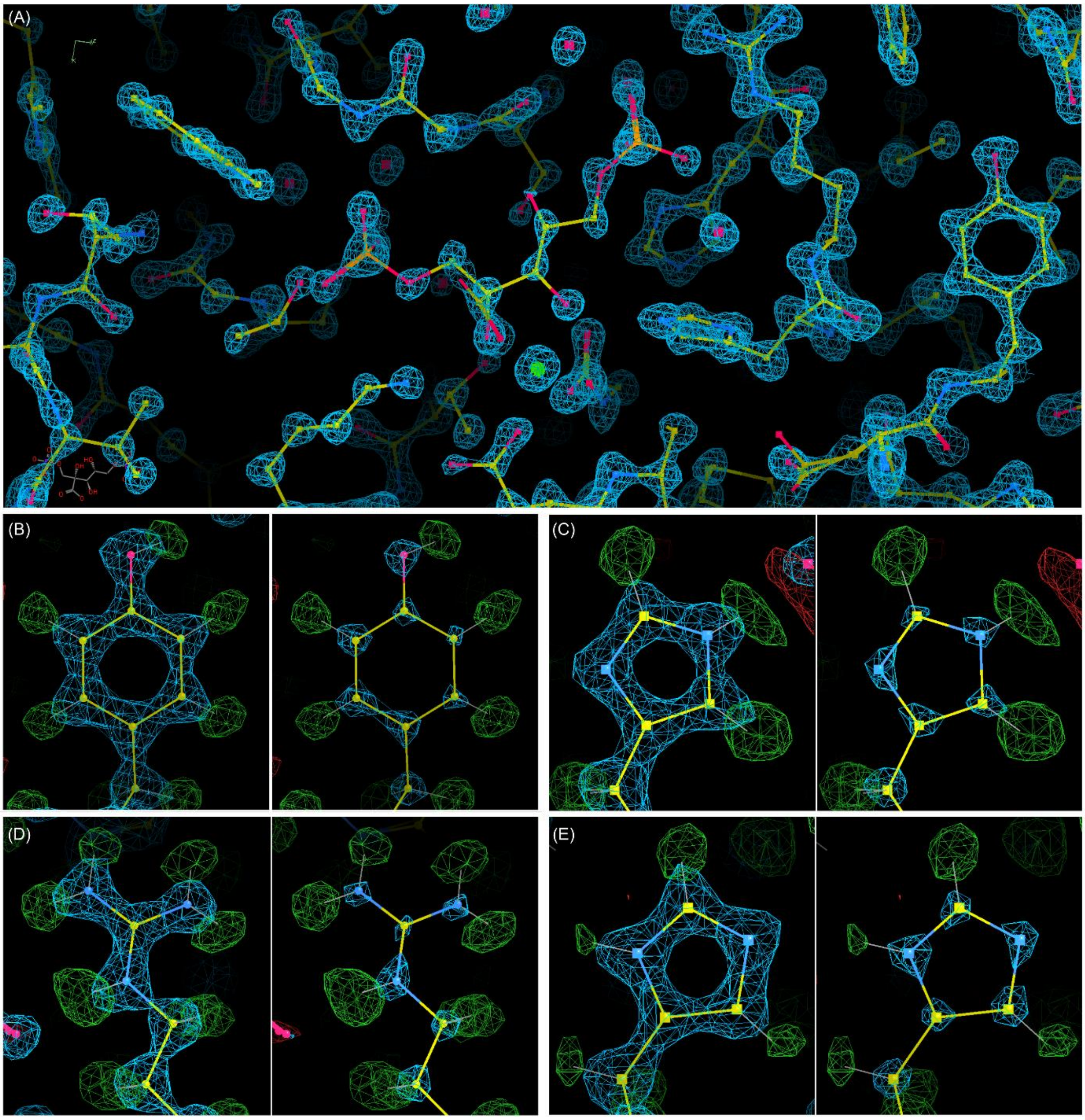

**Supplementary Fig. 1 | Atomic resolution data. (A)** The active site. The $F_o$ map (blue) is displayed at a contour level of 12.6 σ. **(B)** Y269. The $F_o$ maps (blue) are displayed at contour levels of 9.4 σ (left) and 20.3 σ (right). **(C)** H325. The $F_o$ maps (blue) are displayed at contour levels of 9.4 σ (left) and 22.1 σ (right). **(D)** R295. The $F_o$ maps (blue) are displayed at contour levels of 9.4 σ (left) and 22 σ (right). **(E)** H327. The $F_o$ maps (blue) are displayed at contour levels of 9.4 σ (left) and 20.3 σ (right). In (B-E), H-omit $F_o$-$F_c$ maps (green/red) are displayed at a contour level of 10.5 σ. The green densities indicate the positions of hydrogen atoms. All panels were generated with Coot[10] using the consensus structure H-omit mtz file generated by the Servalcat[1] "fofc" command.

### Extended Data

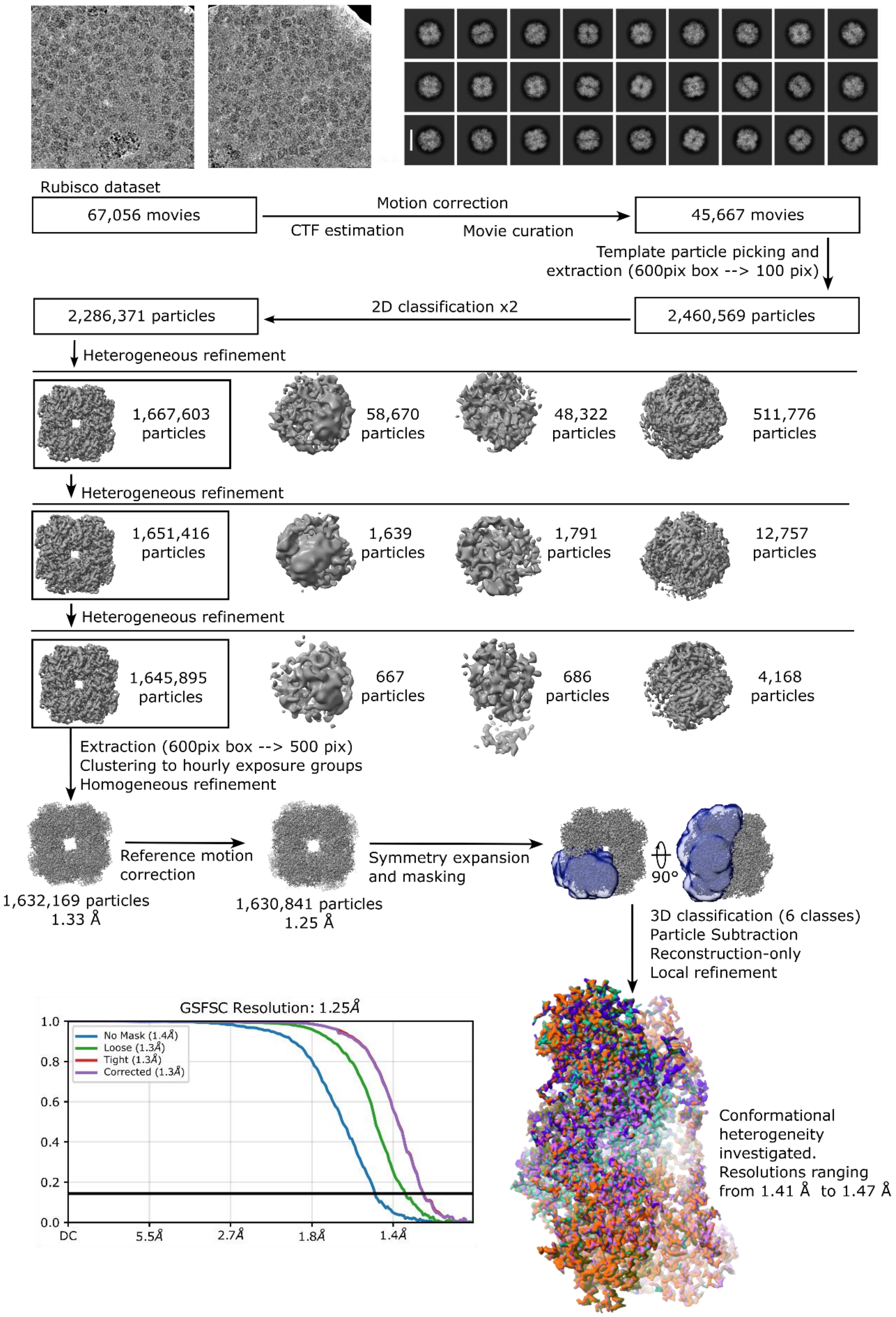

**Supplementary Fig. 2 | Cryo-EM data processing overview.** a. Screening images of rubisco particles on the grid. b. 2D classes. c. Data processing workflow. d. GSFSC curves for the final 1.25 Å reconstruction.

### Extended Data

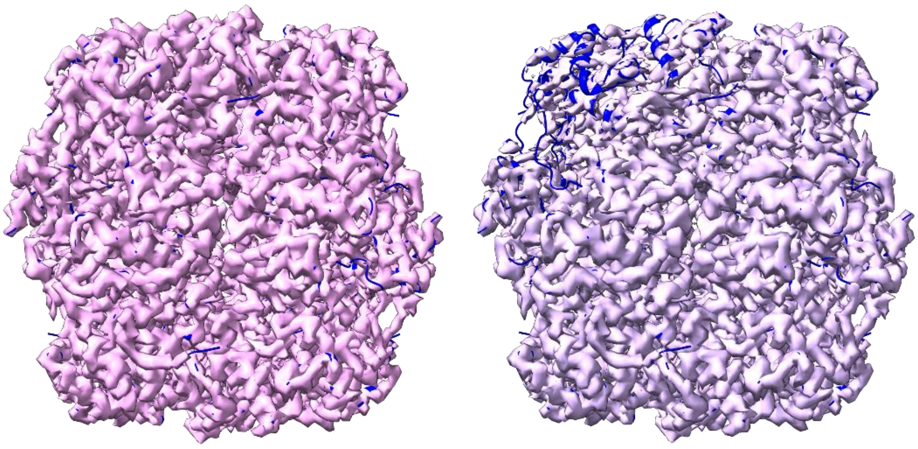

**Supplementary Fig. 3 | Air-water interface damage.** (Left) An undamaged reconstruction. (Right) A reconstruction with air-water interface damage (class 5 in Supplementary Table 2).

### Extended Data

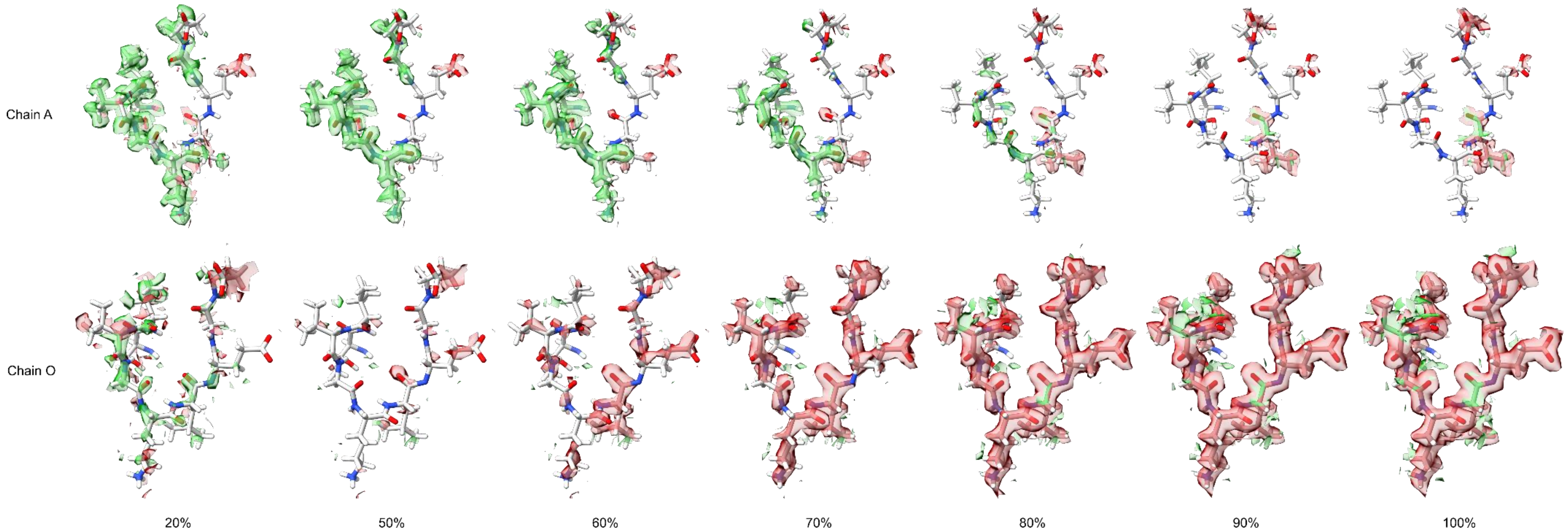

**Supplementary Fig. 4 |Occupancy estimation method**. In all active sites, the occupancy of the N-domain, loop 6, and the C-terminus was estimated (results summarized in Supplementary Table 3) by calculating $F_o$-$F_c$ difference maps with varied occupancy, as shown here for loop 6 in the C-O model. Chain A contains the closed active site, while chain O contains the mixed state active site. One residue in chain A loop 6, L335 has multiple conformations in the closed state. Because the occupancy was set equally to all atoms in loop 6 for the occupancy calculation, the occupancy for L335 is overestimated and negative (red) densities appear over this residue starting at 70% occupancy. The maps are show with absolute map levels of +/- 1.2.

**Extended Data**

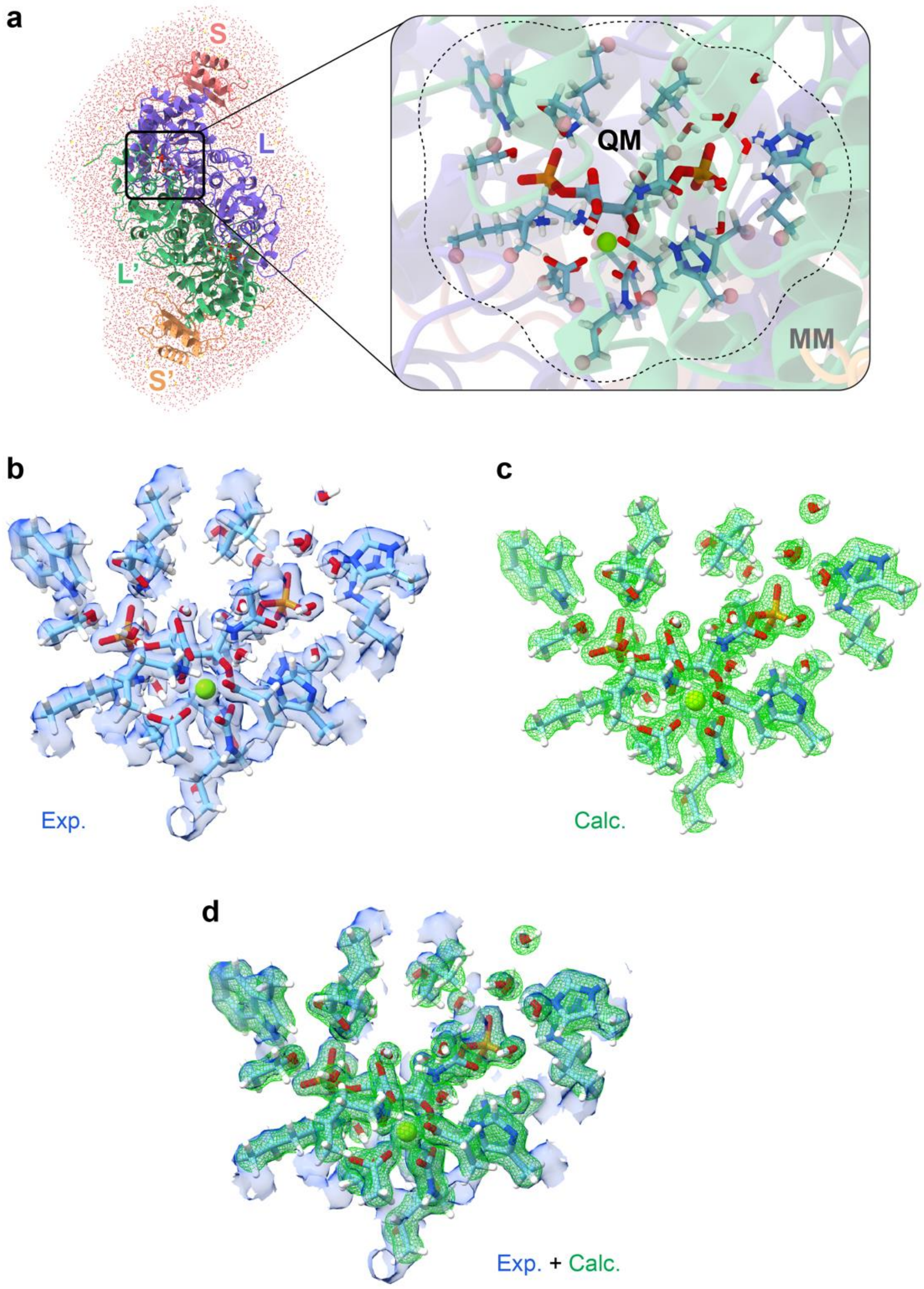

**Supplementary Fig. 5 | QM/MM setup and comparison of cryo-EM and QM/MM ESP maps. a**, Setup of the QM/MM model. **b**, cryo-EM map (Exp.), and **c**, QM/MM electrostatic potential map (Calc.), and the **d**, overlay of the cryo-EM and QM/MM maps.

**Extended Data**

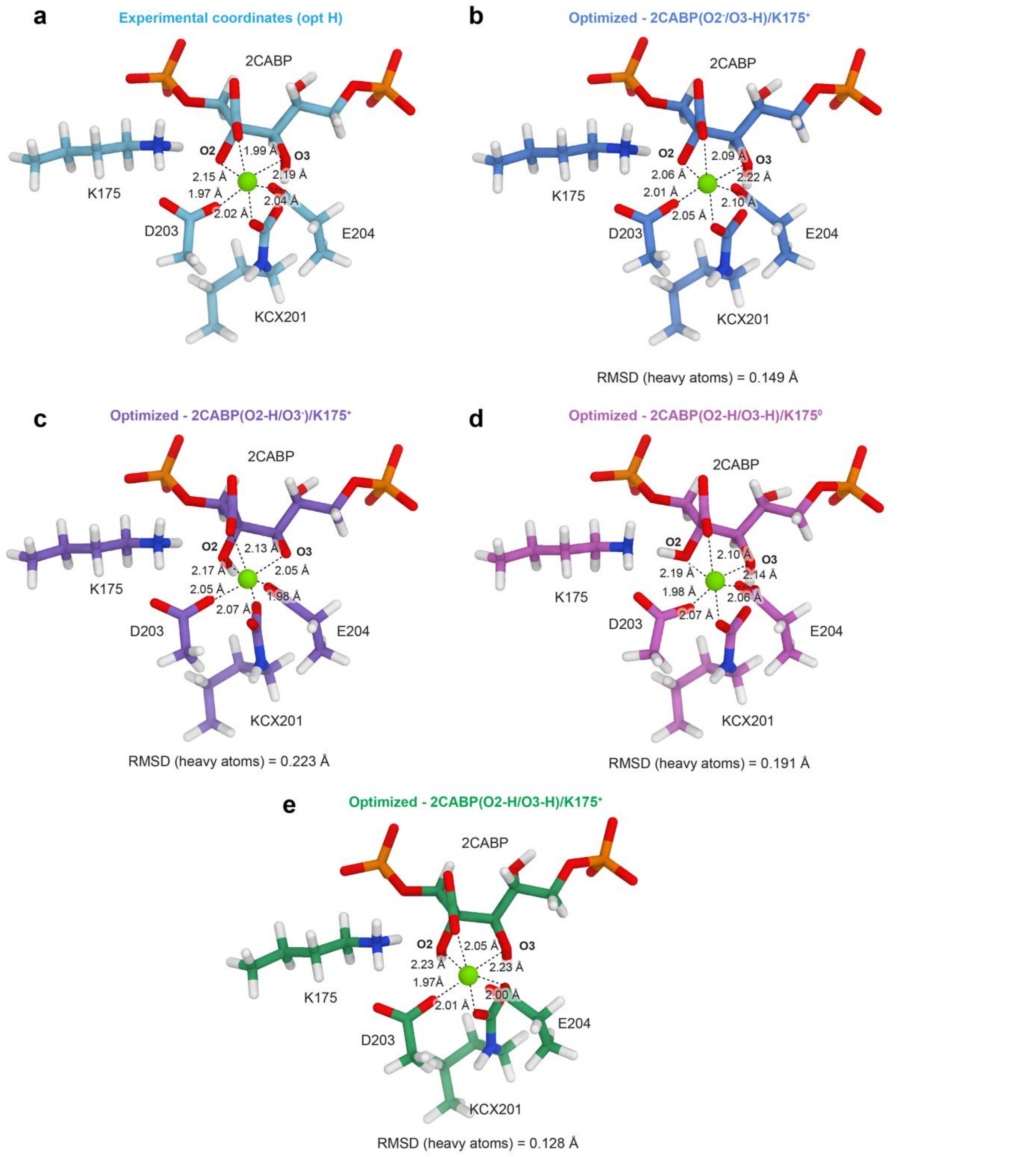

**Supplementary Fig. 6 | QM/MM optimised geometries.** Optimised geometries of the first-metal coordination sphere with different protonation states of the 2CABP (**b-e,** models 4-7, see Supplementary Table 6), compared to **a** (model 1), the coordinates from the cryo-EM map. Relaxed model **5** (**c**) is 3.5 kcal mol$^{-1}$ higher in energy relative to the relaxed model 4 (**b**), whereas relaxed models **4** and **6** (**d**) are isoenergetic (model 6 is 0.06 kcal mol$^{-1}$ higher in energy).

### Extended Data

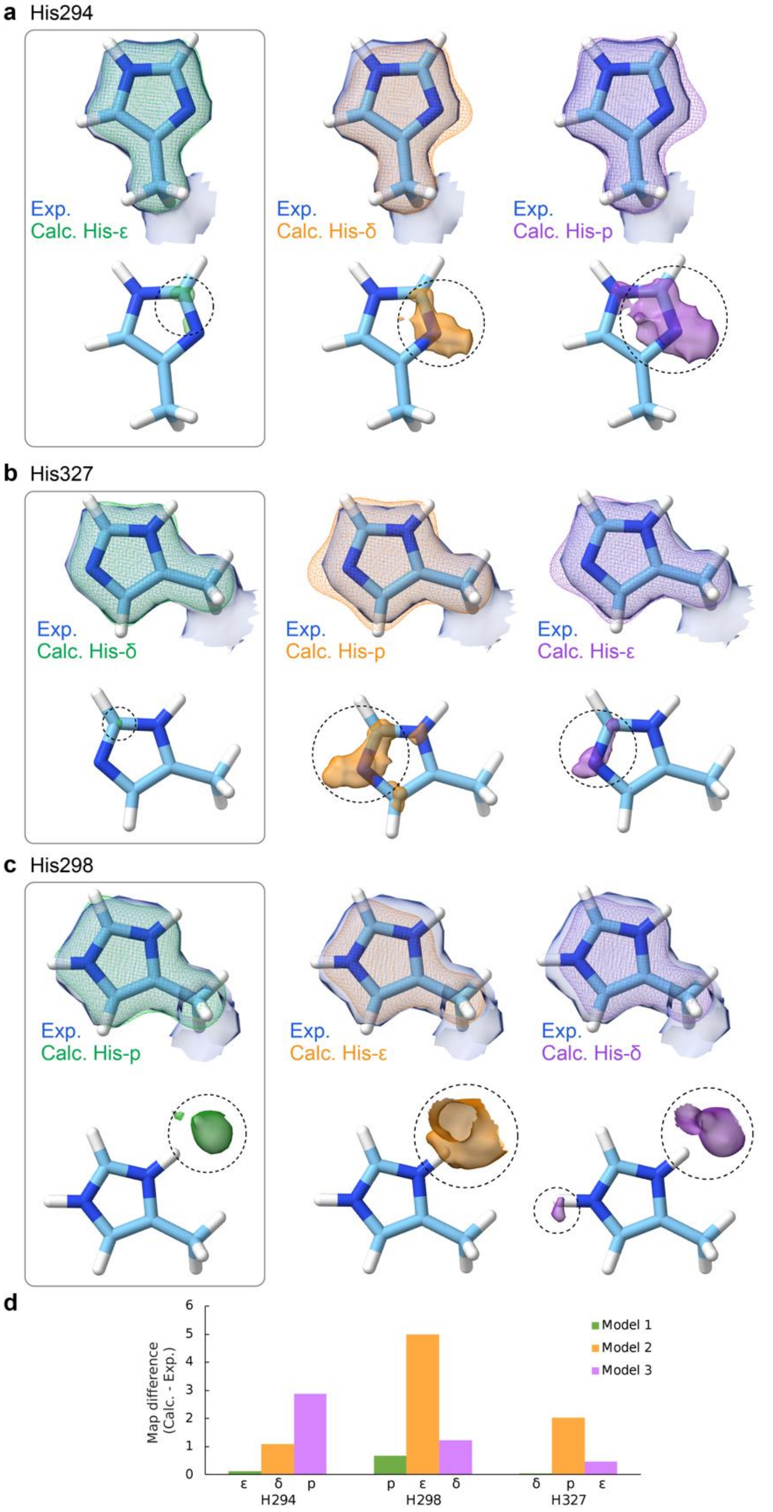

**Supplementary Fig. 7 | Comparison of cryo-EM and QM/MM difference maps.** Comparison of cryo-EM maps (blue transparent surface), and QM/MM electrostatic potential maps (mesh) for different protonation states of the active site histidine residues (Supplementary Table 6: model 1 (green), model 2 (orange), model 3 (purple). **(a)** His294**, (b**) His298 **(c)** His327 (*top:* overlay of cryo-EM map (blue transparent surface, 0.055 absolute map level) and QM/MM-ESP map (coloured mesh, 0.4 absolute map level); *bottom*: difference maps (calculated – experimental) at 0.4 absolute map level. **(d)** Computed volume of the difference maps.

### Extended Data

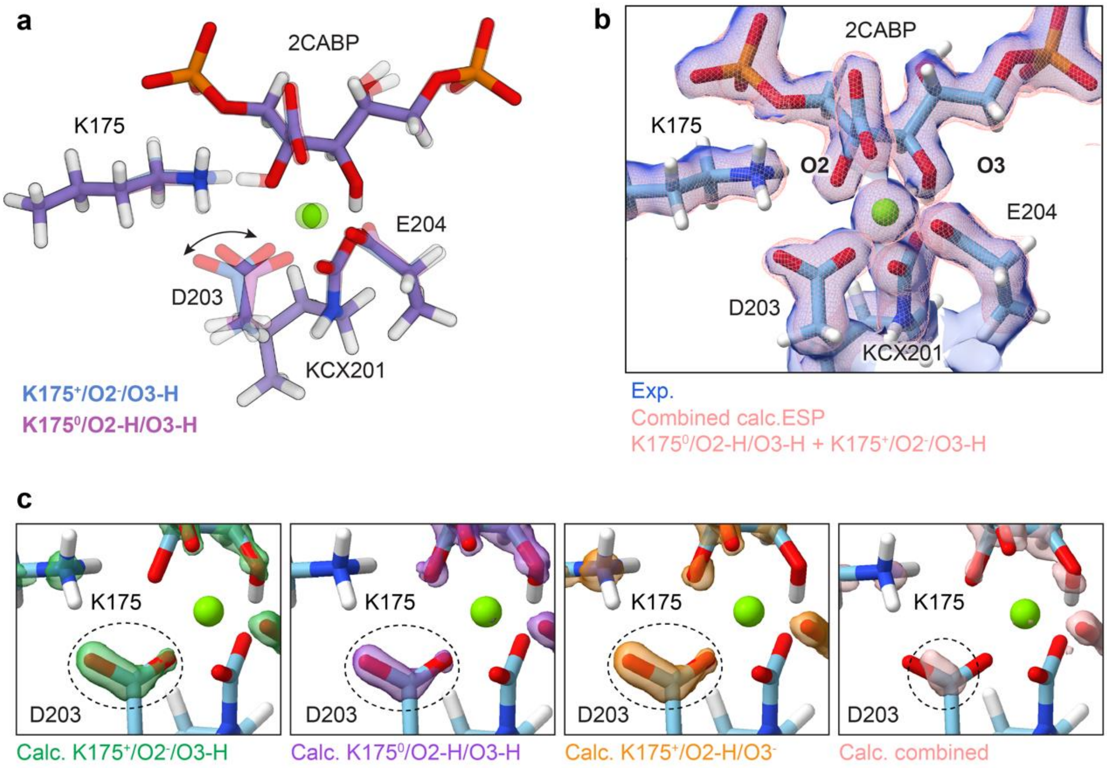

**Supplementary Fig. 8 | Calculated ESP maps from multiple protonation states. (a)** Superposition of optimised geometries with different protonation states of the ligand and protein residues (blue: 2CABP(O2=O$^-$/O3=O-H)/K175$^+$, purple 2CABP(O2=O-H/O3=O-H)/K175$^0$). (**b)** Superposition of the experimental cryo-EM map (blue transparent surface), and the QM/MM-calculated ESP map (pink mesh) as a sum of the ESP maps of the two states represented in a), with equal weight. (**c)** Difference maps between experimental and QM/MM calculated cryo-EM/ESP maps, respectively.

**Extended Data**

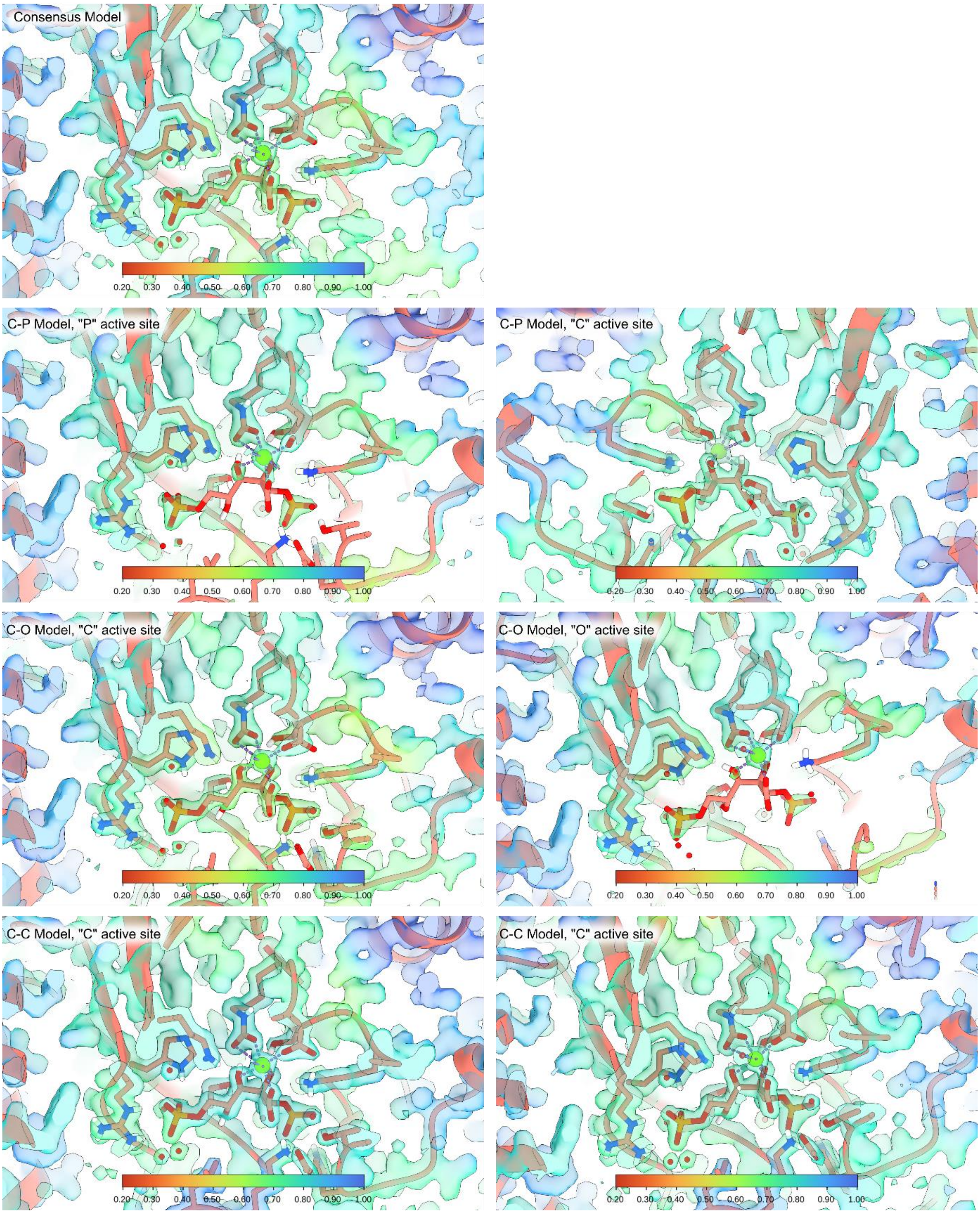

**Supplementary Fig. 9 | OccuPy figures.** The local scale of our cryo-EM maps was calculated with OccuPy[9] for each active site in each model. In the "*P*" and "*O*" active sites, the density in the middle of the 2CABP is very weak, but the phosphates are approximately on the same level as nearby residues, suggesting that the decreased density is the result of flexibility rather than decreased occupancy.

**Extended Data**

**References**

1. Yamashita, K., Palmer, C. M., Burnley, T. & Murshudov, G. N. Cryo-EM single-particle structure refinement and map calculation using Servalcat. *Acta Cryst D* **77**, 1282–1291 (2021).
2. Pettersen, E. F. *et al.* UCSF ChimeraX: Structure visualization for researchers, educators, and developers. *Protein Science* **30**, 70–82 (2021).
3. Nakane, T. *et al.* Single-particle cryo-EM at atomic resolution. *Nature* **587**, 152–156 (2020).
4. Yip, K. M., Fischer, N., Paknia, E., Chari, A. & Stark, H. Atomic-resolution protein structure determination by cryo-EM. *Nature* **587**, 157–161 (2020).
5. Agirre, J. *et al.* The CCP4 suite: integrative software for macromolecular crystallography. *Acta Cryst D* **79**, 449–461 (2023).
6. Pierce, J., Tolbert, N. E. & Barker, R. Interaction of ribulosebisphosphate carboxylase/oxygenase with transition-state analogs. *Biochemistry* **19**, 934–942 (1980).
7. Jordan, D. B. & Ogren, W. L. The CO2/O2 specificity of ribulose 1,5-bisphosphate carboxylase/oxygenase. *Planta* **161**, 308–313 (1984).
8. Schloss, J. V. Comparative affinities of the epimeric reaction-intermediate analogs 2- and 4-carboxy-D-arabinitol 1,5-bisphosphate for spinach ribulose 1,5-bisphosphate carboxylase. *Journal of Biological Chemistry* **263**, 4145–4150 (1988).
9. Forsberg, B. O., Shah, P. N. M. & Burt, A. A robust normalized local filter to estimate compositional heterogeneity directly from cryo-EM maps. *Nat Commun* **14**, 5802 (2023).
10. Emsley, P., Lohkamp, B., Scott, W. G. & Cowtan, K. Features and development of Coot. *Acta Cryst D* **66**, 486–501 (2010).
